# Coupling phenotype stability to growth rate overcomes limitations of bet-hedging strategies

**DOI:** 10.1101/2022.04.12.488059

**Authors:** Daan H. de Groot, Age J. Tjalma, Frank J. Bruggeman, Erik van Nimwegen

## Abstract

Microbes in the wild face highly variable and unpredictable environments, and are naturally selected for their average growth rate across environments. Apart from using sensory-regulatory systems to adapt in a targeted manner to changing environments, microbes employ bet-hedging strategies where cells in an isogenic population switch stochastically between alternative phenotypes. Yet, bet-hedging suffers from a fundamental trade-off: increasing the phenotype switching rate increases the rate at which maladapted cells explore alternative phenotypes, but also increases the rate at which cells switch out of a well-adapted state. Consequently, it is currently believed that bet-hedging strategies are only effective when the number of possible phenotypes is limited and when environments last for sufficiently many generations. However, recent experimental results show that gene expression noise generally decreases with growth rate, suggesting that phenotype switching rates may systematically decrease with growth rate. We here show that such growth rate dependent stability (GRDS) can almost completely overcome the trade-off that limits bet-hedging, allowing for effective adaptation even when environments are diverse and change rapidly. GRDS allows cells to be more explorative when maladapted, and more phenotypically stable when well-adapted. We further show that even a small decrease in switching rates of faster growing phenotypes can substantially increase long-term fitness of bet-hedging strategies. Together, our results suggest that stochastic strategies may play an even bigger role for microbial adaptation than hitherto appreciated.

## Introduction

Many microbial organisms exhibit a remarkable ability to adapt their internal state to environments that are highly variable and can change in unpredictable ways. For example, not only will there be different types of carbon sources, nitrogen sources, and amino acids available in different environments, also the concentrations of all these nutrients may vary over orders of magnitude. In addition, general variables such as temperature, pH, osmotic pressure, and oxygen availability will vary, and cells may have to withstand a wide array of specific stresses such as antibiotics or reactive oxygen species. It is remarkable that microbes appear able to adapt to this enormous number of possible combinations of environmental variables, not only because it requires coordinating the expression of many genes, but also because it seems unlikely that the microbes can have been specifically selected for adapting to all these environments. For example, *E. coli* is able to adapt its gene expression in order to allow it to grow in fully deuterated water, a highly unnatural condition that it almost certainly never encountered in the wild [1].

It is well-known that microbes have evolved sensory-regulatory machinery that can sense a large variety of internal and environmental variables, and adapt gene expression patterns in response. In principle, the more information an organism gathers about its environment, the better it can adapt to it [2,3]. However, sensory-regulatory strategies face a number of limitations. First, sensing is limited to those environmental variables for which the organism has evolved sensors, which is likely only a subset of the many additional environmental variables that affect optimal gene expression states. Second, given the small number of molecules involved, there are fundamental thermodynamic limits on the accuracy with which cells can gather information about their environment [4,5]. Third, even if cells are able to gather accurate information about the state of environmental variables, the processing and integrating of this information so as to optimally set gene expression levels is a non-trivial regulatory problem, and the regulation of gene expression is itself also significantly affected by thermodynamic noise. Finally, there may be intrinsic costs associated with sensory-regulatory machinery, be it through the cost of expressing proteins that do not directly contribute to growth [6,7], or due to energetic costs [8].

Instead of adapting gene expression in a regulated manner, microbial populations can also adapt to changing environments by using a so called bet-hedging strategy [9–15]. Isogenic cells then explore alternative phenotypes by switching stochastically between different phenotypes, and the subpopulation with a fast-growing phenotype is automatically expanded because of its higher growth rate. This strategy allows microbes to adapt to a wide variety of unexpected environmental changes, including those that they are unable to sense.

However, the long-term fitness that can be attained with this bet-hedging strategy, i.e. the long-term average population growth rate [16,17], is limited by an intrinsic trade-off: increasing the stochastic phenotype switching rate speeds up adaptation to new environments, but it also decreases the long-term growth rate in each environment since it increases the rate at which cells switch out of well-adapted phenotypes [18]. Due to this inherent trade-off, as demonstrated both by theoretical modelling [10,18,19] and by experimental approaches [20,21], bet-hedging strategies are only effective when durations of environments are large relative to the doubling times of the cells, and when the number of possible environments that populations need to anticipate is limited.

Several recent studies have observed that gene expression noise generally increases at low growth rates [22–24]. Since many phenotype switches may be driven by gene expression fluctuations [25–29], these observations suggest that there may be an intrinsic coupling between the growth rate of cells and their phenotypic stability. That is, stochastic phenotype switching rates may naturally be higher for slow growing cells than for fast growing cells. Intuitively, it seems that such Growth Rate Dependent Stability (GRDS) could benefit bet-hedging strategies since it would reduce the rate at which well-adapted cells switch to maladapted phenotypes, while at the same time increasing the rate at which maladapted cells explore alternatives. Indeed, as has been shown by Kaneko and coworkers in a non-evolutionary setting [30,31], when phenotypic stability increases with growth rate, the distribution of phenotypes in the population is shifted toward faster growing phenotypes.

Here we systematically investigate the effect of GRDS on the performance of bet-hedging. In particular, we extend the basic model of a population evolving in a changing environment introduced in [18], and using a combination of analytical solutions, mathematical proofs and simulations, we determine how GRDS affects the long-term average growth rates that stochastically switching populations can achieve. We show that even a small growth rate dependence of the phenotype switching rates immediately increases the average growth rate of bet-hedging populations, and that GRDS can completely resolve the inherent trade-off of traditional bet-hedging strategies when the ratio of switching rates of slow and fast growing cells is sufficiently high. We also find that GRDS improves average population growth rates through two qualitatively distinct mechanisms depending on whether environment durations are short or long relative to the doubling time of adapted phenotypes. Taken together, our results show that GRDS dramatically expands the range of scenarios for which stochastic bet-hedging strategies can attain high long-term average growth rate.

## Results

### Model setup

To investigate the effects of growth rate dependent stability (GRDS) on bet-hedging strategies for microbial populations growing in changing environments we extend the general model introduced by Kussell and Leibler [18]. We consider a population of cells that switch stochastically between *n* + 1 discrete phenotypes and grow in an environment that switches stochastically between *m* discrete environment-types. For a given realisation of the stochastic environment switches, the sequence of environments is described by the function 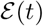, denoting which environment-type is present at time *t*. The average time that an environment of type *J* remains before it switches to another environment-type is *τ_J_*. The order in which environment-types occur is determined by a Markov chain: upon a switch, the probability that environment *J* is followed by environment *I* is denoted by *b_IJ_*, where ∑*_I_ b_IJ_* = 1 and *b_II_* = 0. In environment *J*, cells with phenotype *i* grow at a rate 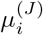, and switch stochastically from phenotype *j* to *i* at rate 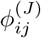.

The population is assumed to be sufficiently large such that the population dynamics can be modelled by a set of deterministic differential equations. The state of the population is described by an (*n* + 1)-dimensional vector ***s***, containing the number of cells of each phenotype, whose dynamics are described by

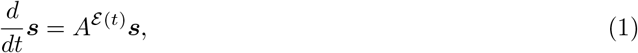

where 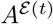 is the time-evolution matrix of environment 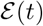. The components 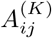 of the time-evolution matrix of environment *K* are given by

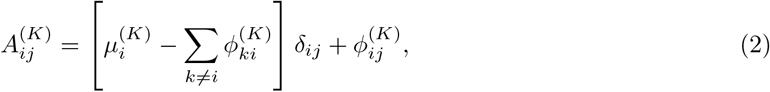

where *δ_ij_* is the Kronecker delta-function that is one when *i* = *j* and zero otherwise.

Up to this point, this model is identical to the general model used in [18] for modelling a bet-hedging population in fluctuating environments. In the classical bet-hedging scenario of [18], the switching rates 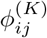 are assumed independent of the environment, i.e. 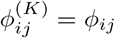 for all *K*. To investigate the effects of GRDS, we extend this classical model by allowing the switching rates 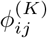 to be functions of the current growth rate of the cell, i.e. 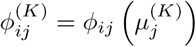, and generally assume that *ϕ_ij_*(*μ*) is a decreasing function of the growth rate *μ*. We will quantify the strength of GRDS by the parameter *r*, which is the ratio of switching rates between the fastest and slowest growing phenotypes, i.e. when phenotype *j* achieves its maximal growth rate in environment *J* and its minimal growth rate in environment I we get 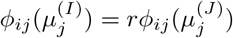. Note that the case where *r* =1 thus corresponds to classical bet-hedging without GRDS.

The total number of cells at time t is given by *N*(*t*) = ∑_*i*_ *s_i_*(*t*) and we are generally interested in the average growth rate *G* of the population over a long sequence of environments, i.e.

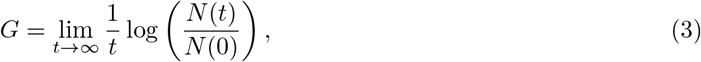

which quantifies the ‘fitness’ of a given strategy [3,16,17]. In particular, we will compare the long-term growth rates *G* that are obtained with classical bet-hedging with those obtained with GRDS.

### A toy example that qualitatively illustrates the benefits of GRDS

We use a toy example of this general model to illustrate the differences in behaviour between classical bet-hedging and bet-hedging with GRDS (Fig. 1). In this example there are are only 3 environments and phenotypes (shown as purple, red, and green); in each environment, one phenotype is optimal and leads to a growth rate of *μ*_1_ = 1.0, while the other phenotypes have a growth rate of *μ*_0_ = 0. For the population dynamics shown here, all cells start out in the green phenotype and encounter the purple environment, followed by the red environment; both environments have a duration of *T* = 10. We assume that cells in the optimal phenotype switch with a rate *ϕ*, while the other cells switch at a rate *rϕ* where *r* = 10 for bet-hedging with GRDS (and *r* = 1 for classical bet-hedging). The switching rate *ϕ* was optimised to maximise *G* for both strategies separately.

**Figure 1.**
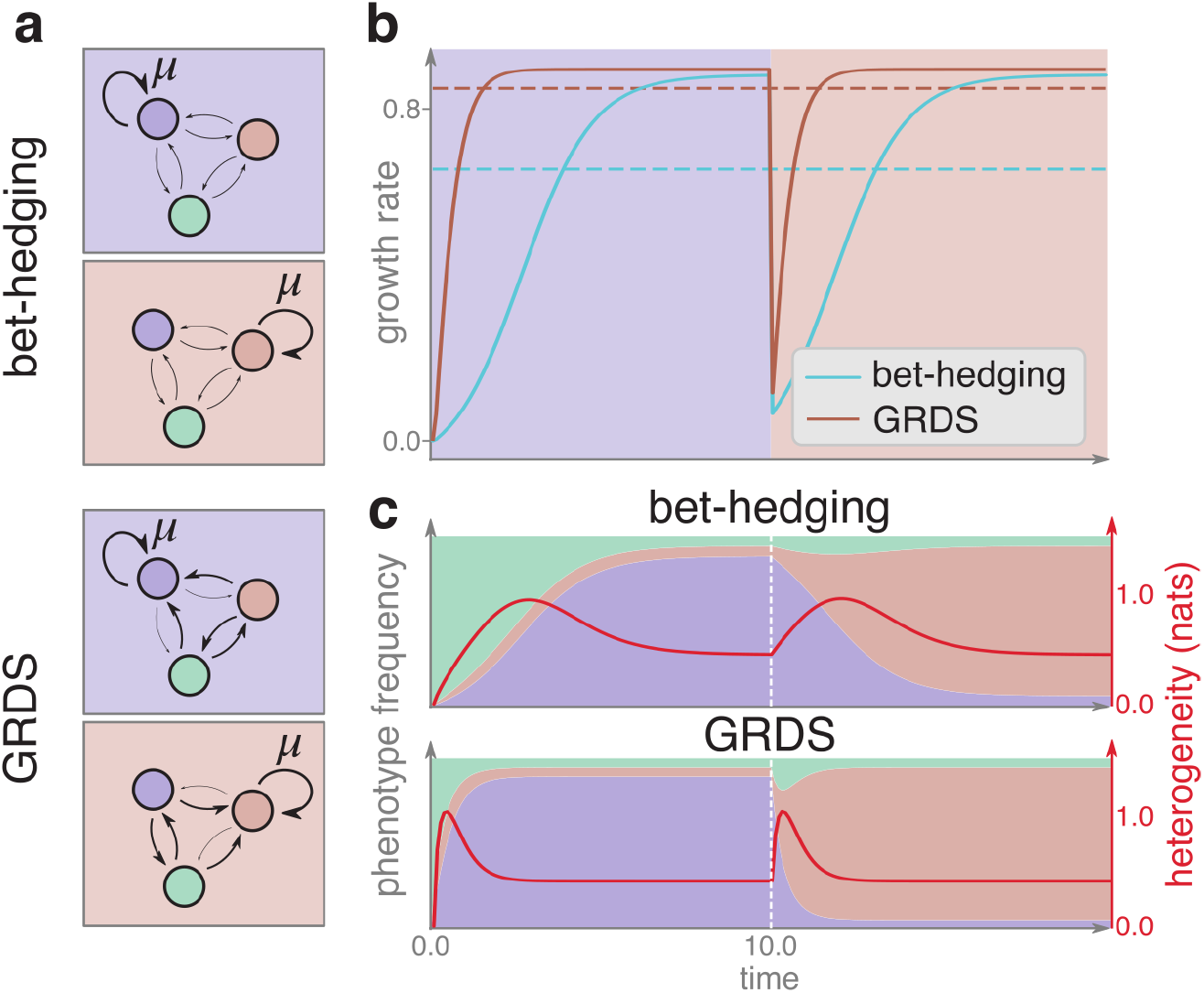
A toy example illustrates how GRDS increases the effectiveness of bet-hedging. **a)** Growth was simulated in a sequence of two environments (purple and red background) for cells with three different phenotypes: optimal for either the purple, red or green environment. The growth of a phenotype is indicated by the arrow marked by *μ*, while the other arrows indicate switching between phenotypes. Bolder lines indicate higher rates. Random phenotype switching rates are either constant (bet-hedging) or growth rate dependent (GRDS). The optimal tuning of the switching rates *ϕ* to maximise the average growth rates G resulted in *ϕ* = 0.13 for bet-hedging without GRDS, and *ϕ* = 0.26 for bet-hedging with GRDS. **b)** Population growth rates as a function of time for bet-hedging without GRDS (blue curve) and with GRDS (red curve), starting from an initial condition with all cells in the green phenotype. The average growth rates for both strategies are indicated by dashed lines. **c)** Time courses of the fractions of the population in each of the phenotypes (colors) for the bet-hedging (top panel) and GRDS (bottom panel) strategies. The red curves show time courses of the population heterogeneity, defined as the entropy of the distribution of phenotypes in the population. The parameters used for these simulations, as described in the Model setup-section, were *n* + 1 = 3, *T* = 10, *μ* = 1.0 and *r* = 10.

The inherent trade-off of classical bet-hedging is illustrated in Figure 1a. Because the switching rate is independent of the environment, a cell in a given phenotype is equally likely to switch independent of whether this phenotype is well or badly adapted to the current environment. Thus, increasing the switching rate will not only increase the rate at which cells in maladapted phenotypes explore alternative phenotypes, it will also increase the rate at which cells with optimal phenotype switch to worse phenotypes. The trade-off thus arises because a high long-term growth rate requires a relatively low switching rate, while fast adaptation requires a high switching rate. Indeed, at the switching rate *ϕ* that optimises this trade-off, the adaptation takes about half the environment duration, but speeding this up would require a higher switching rate which would decrease the long-term population growth rate (Fig. 1b, blue curve).

GRDS can largely resolve this trade-off by decoupling the rate of exploration by non-adapted cells, i.e. *rϕ*, from the rate of switching of well-adapted cells, i.e. *ϕ*. This makes it possible to speed up the adaptation to a new environment, while the same long-term population growth rate is reached (Fig. 1b, red curve). Note that the population in each environment stabilises with a similar fraction of cells in suboptimal phenotypes as with the classical bet-hedging strategy (Fig. 1c, bottom panel). This shows that, at least in this setting, GRDS mainly works because it allows for a ‘panic mode’: immediately after an environment change the growth rate of the majority of cells drops so that their phenotype switching rates increase, quickly generating a heterogeneous population of cells that explore different phenotypes. Moreover, these exploring cells only stabilise once they find a phenotype that supports fast growth. Since these adapted cells are both more stable and grow faster, the population quickly becomes dominated by optimised cells again (Fig. 1c, bottom panel).

### A minimal model of bet-hedging with GRDS

To quantify the effect of GRDS on bet-hedging strategies we first analyse a model in which the number of parameters is reduced to a minimum and which can be solved analytically. In this minimal model we set the number of environments and phenotypes both to *n* + 1. We assume each environment lasts for a fixed time T and then switches to another environment with uniform probability, i.e. *b_IJ_* = 1/*n* for all *I* ≠ *J*. In addition, we assume that there are only two possible growth rates, a ‘fast’ growth rate *μ*_1_ and a ‘slow’ growth rate *μ*_0_, i.e. *μ*_1_ > *μ*_0_, and that in each environment *I*, only cells in a ‘good’ phenotype *i* = *I* grow at the fast rate *μ*_1_ and cells in the *n* (‘bad’) other phenotypes grow at the slow rate *μ*_0_. Regarding the phenotype switching, we assume that whenever a cell switches its phenotype, it is equally likely to switch to any of the *n* other phenotypes. Finally, to tune the overall switching rates and amount of GRDS we assume that cells in the fast growing phenotype *i* = *I* switch out of their phenotype at a total rate *ϕ*, whereas cells in any of the *n* slow growing phenotypes switch out of their phenotype at a total rate *rϕ* with *r* ≥ 1. Thus, formally the switching rates are given by 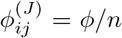 for all *i* ≠ *j* when *j* = *J*, and 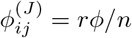 for all *i* ≠ *j* and *j* ≠ *J*.

Because switching rates to and from all bad phenotypes are equal in this model, we can describe the state of the population by the number of cells *s_g_*(*t*) and *s_b_*(*t*) in the good and bad phenotypes, respectively (see SI 1.B.1 for more details). Without loss of generality we can restrict ourselves to solving models where the bad phenotype does not grow at all, i.e. *μ*_0_ = 0, and measure time in units such that *μ*_1_ = 1 (see SI Section 1.B). Solutions for any other setting of the growth rates *μ*_1_ and *μ*_0_ can then be obtained by rescaling and shifting the resulting time dynamics. This leaves us with the following differential equations for the number of cells with good and bad phenotypes

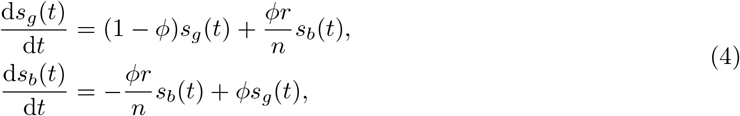

where *ϕ* is the switching rate of adapted (‘good’) phenotype cells and *rϕ* the switching rate of cells in a bad phenotype. Note that, because a cell in a bad phenotype can switch to *n* other phenotypes and only one of these is the good phenotype, the effective switching rate of cells from a bad to the good phenotype is *ϕr/n*. The equations (4) would not change if instead of one good phenotype and *n* bad ones, we assumed there were *K* good phenotypes and *nK* bad ones. This suggests interpreting the probability 1/*n* more generally as the probability that, under a stochastic phenotype switch, a cell in a bad phenotype will switch to a good phenotype. That is, the parameter *n* quantifies how rare fast growing phenotypes are and thereby quantifies the complexity of the fluctuating environment.

When the environment changes, a new phenotype will become optimal so that all cells that were in the good phenotype now find themselves in a bad phenotype, whereas some cells that were in a bad phenotype may coincidentally find themselves in the new good phenotype. In our analysis of the minimal model, we will make the approximation that all cells that were in a bad phenotype in the previous environment have a probability 1/*n* to find themselves in the good phenotype of the new environment. Thus, if we denote the fraction of adapted cells at the end of an environment as *p(T*), the initial fraction of adapted cells in the next environment is (1 – *p*(*T*))/*n*. In Section SI 1.C.2 we explain why, under mild conditions on the model parameters, this approximation is justified.

The dynamics of this system can be solved analytically (see SI 1.C and 1.D) yielding an expression for the long-term average growth rate G that is fully determined by the four parameters *T, n, r* and *ϕ*. Of these parameters, *n* and *T* parametrise the regulatory problem that the microbial population faces: the complexity of the environment is set by *n*, and *T* sets the number of generations between environmental changes which determines the relative importance of fast adaptation versus a high stationary growth rate. In turn, the strength of GRDS r and the switching rate *ϕ* set the behaviour of the cellular population.

To further analyse this minimal model, we assume that natural selection has optimised the switching rate *ϕ* to maximise the average growth rate *G*. In Figure 2 we systematically investigate how the resulting average growth rate *G* varies with *n, T*, and the strength of GRDS *r*. We vary *n* and *T* on the vertical axes of Figures 2a-b, while the strength of GRDS increases along the horizontal axes. The growth rates for classical bet-hedging correspond to *r* = 1 and are thus shown along the vertical axes in these plots. We see that, as derived previously [18], the fitness of a bet-hedging population decreases with the complexity of the environment *n* and increases with the environment duration *T* for classical bet hedging. Increasing GRDS by moving away from the vertical axes at *r* = 1 in Figs. 2a-b, we see that GRDS can dramatically increase the average growth rate and that increasing *r* is always beneficial. That is, at least in this minimal model, evolution would favour making the ratio *r* between the switching rates at slow and fast growth as large as possible. Moreover, provided the strength of GRDS *r* is made sufficiently large, the average growth rate *G* can approach the theoretical optimum *G* = 1 arbitrarily closely.

**Figure 2.**
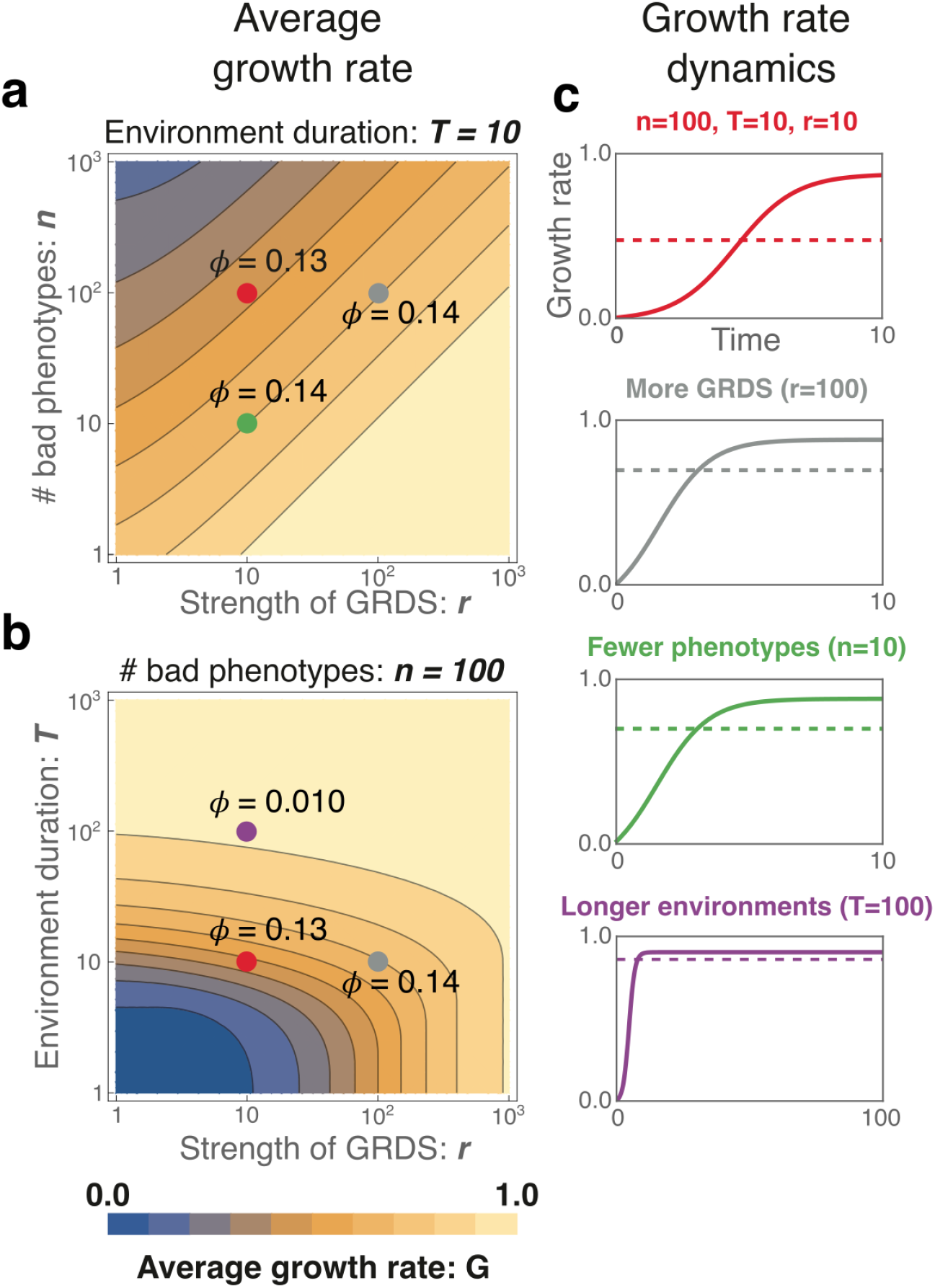
Effect of GRDS on the average growth rate of bet-hedging strategies as a function of environment complexity and duration. The contourplots in panels **a** and **b** show the average population growth rate (*G*) at optimal switching rate *ϕ* as a function of the strength of GRDS (horizontal axes, both panels) and either the number of environments (**a**) or the environment duration (**b**). Since *μ*_1_ = 1 and *μ_0_* = 0, 0 ≤ *G* ≤ 1 and the contours occur at integer multiples of 0.1. The optimal switching rates *ϕ* at 4 example parameter settings (colored dots) are indicated in the contour plots. All axes are on logarithmic scales. Contour plots for additional parameter settings are shown in Suppl. Figs. S3 and S4. **c**: Population growth rate versus time over the course of one environment for the parameter sets indicated by coloured dots in panels **a** and **b**. The dashed lines indicate the average growth rate *G*. Note that the grey, green and purple parameter settings each differ from the red parameter setting by a change in one parameter.

To illustrate the dependence of the average growth rate *G* on the parameters, we start from a reference parameter set *n* = 100, *T* = 10, *r* = 10 (red dots in Fig. 2a-b) and either increase *r* by a factor of ten (grey dots Fig. 2a-b), decrease *n* by a factor ten (green dot in Fig. 2a) or increase *T* by a factor ten (purple dot in Fig. 2b). When the strength of GRDS *r* is increased by a factor ten, this has a similar effect on growth rate *G* as decreasing the number of bad phenotypes *n* by ten, and more generally the approximately straight diagonal contours in Fig. 2a show that *G* effectively only depends on the ratio *r/n.* This can be understood by noting that the population dynamics of (4) depends on *r* and *n* only through the ratio *r/n*. Although the initial fraction of good phenotype cells after an environment switch does depend directly on *n* and not *r*, we find that this is negligible when *T* is not small.

#### Long environment durations

Of the three parameter changes in Fig. 2, the optimal switching rate *ϕ*^opt^ only changes significantly with the change in environment duration *T*, and is largely insensitive to changes in the other parameters. Indeed, as shown in Suppl. Fig. S7, we find that when T is sufficiently large (i.e. around *T* = 10 generations or larger), and *r* is not too large (i.e. *r* < *nT*, as discussed below), the optimal switching rate is just the inverse of the environment duration:

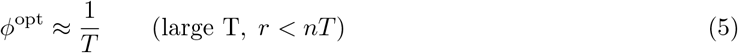

and in SI Section 2.B.2 we mathematically prove that this relationship holds exactly in the limit of long environment durations. Thus, the optimal phenotype switching rate for cells in the fast growth phenotype exactly equals the rate at which the environment changes. This extends a classical result for conventional bet-hedging which states that the probability of using a strategy should equal the probability that the strategy will become useful in the future [18,19,32,33]. Note that, in this large *T* parameter regime, the optimal switching rate is independent of *r* and thus the same for classical bet hedging and bet hedging with GRDS. This shows that the benefits of GRDS derive not from a greater stability of cells in the optimal phenotype but from the increased switching rate of the slow growing cells. That is, GRDS allows slow growing cells to ‘panic’ and rapidly explore different phenotypes until a fast growing phenotype is found. Such an adaptive transition of the population from stable to explorative and back is impossible without GRDS.

We also derived an analytical approximation for *G* in the parameter regime where environment durations are sufficiently long for the population to reach its steady-state distribution of phenotypes (see SI 2.B.2):

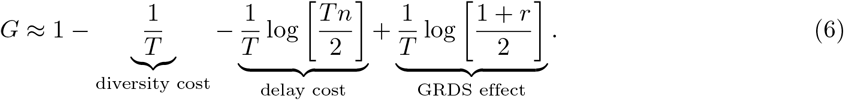

The terms in this equation have intuitive interpretations: first, at the optimal switching rate *ϕ*^opt^ = 1/*T*, the long-term growth rate is 1 – 1/*T* because a fraction 1/*T* of the population will be in non-growing phenotypes. In [18] this is referred to as the ‘diversity cost’ of bet-hedging. Second, at the end of a given environment, the non-growing cells will be equally distributed over the *n* non-growing phenotypes, so that the fraction of cells in the ‘good’ phenotype just after an environment switch will be 1/(*nT*). The ‘delay cost’ log[*nT*/2]/*T* corresponds to the reduction of the average growth rate from having to expand this small subpopulation. The final term log[(1 + *r*)/2]/*T* is unique to bet-hedging with GRDS and quantifies the extent to which GRDS can compensate the intrinsic delay and diversity costs of bet-hedging. We have validated numerically that (6) provides an excellent approximation to *G* as long as environment durations are not short, i.e. *T* ≥ 10 generations (Suppl. Fig. S8). Supplemental Fig. S8 also shows that once *r* becomes so large that the delay costs are fully compensated, i.e. when *r* ≥ *Tn*, Equation (6) starts to overestimate *G*, and the true growth rate saturates towards *G* = 1 when *r* increases further. As discussed in the next section, in this regime the optimal switching rate no longer equals 1/*T* and the approximation breaks down.

We derived an analogous expression to Equation (6) for the more general model, including differing average environment durations *τ_J_* and transition probabilities *b_IJ_*, using the same assumptions as were used in [18] for the case without GRDS (see SI 2.B.2). In this more general setting GRDS still increases the average growth rate with log[(1 + *r*)/2]/*T*, giving

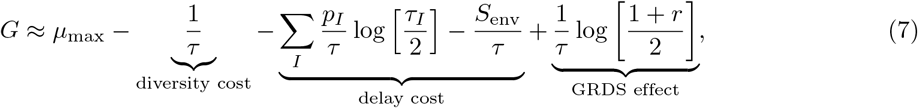

where *μ*_max_ is the growth rate of the adapted phenotype in each environment, *τ* = ∑_*I*_ *p_I_τ_I_* is the average environment duration, *p_I_* is the probability of environment *I* occurring, and *S*_env_ = = –∑_*I,J*_ *b_I,J_p_J_* log(*b_IJ_*) is the environment entropy, which quantifies how hard it is on average to predict the next environment given the current environment. In Equation (6), this entropy term was log(*n*) because we assumed that after each environment all other environments were equally likely to occur next. In general, the change of environments can be much more predictable, for example when environment *I* is always followed by environment *J* (i.e. *b_IJ_* = 1), which decreases the delay cost for bet-hedging since switching rates can be adapted to this predictability.

Equation (7) shows that GRDS can compensate the intrinsic costs of bet-hedging, including the uncertainty about the environment captured by *S*_env_, revealing that GRDS makes cells effectively learn about their environment through their growth rate. The logarithmic dependence also shows that even a slight growth rate dependence of the phenotype switching rates can already provide a substantial fitness advantage. Finally, Equation (7) implies that a population employing GRDS outgrows a population employing classical bet-hedging by a factor (1 + *r*)/2 over the course of each environment, independent of the number of environment-types, the transition rates *b_IJ_* between them, or their durations *τ_I_*.

#### Short environment durations

For classical bet-hedging it is known that, when environment durations are short relative to the doubling time of the fastest growing cells, bet-hedging is ineffective because natural selection has no time to expand the subpopulation of fast growing cells. However, Fig. 2b shows that, even when *T* = 1, bet-hedging with GRDS can reach close to maximal fitness provided that *r* is made sufficiently large. Interestingly, in this small *T* regime, the optimal strategy is to switch as fast as possible, i.e. *ϕ*^opt^ → ∞. In this limit of very fast switching, the steady-state fraction of cells in the fast growth phenotype is given by *P*_*ϕ*→∞_ = *r*/(*n* + *r*). Due to the fast switching this steady-state is reached very quickly and, consequently, the average population growth rate becomes

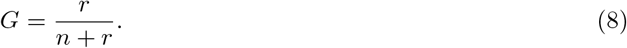

Although an infinite switching rate is not realistic, this strategy of fast switching is already effective when *ϕ* is large compared to the rate at which environments switch (1/*T*) and compared to the growth rate of adapted cells (*μ* = 1).

GRDS can thus aid adaptation through two qualitatively different strategies. When environment durations *T* are large, fast growing cells come to dominate the population by outgrowing the slow growing cells, and the optimal strategy is for fast growing cells to switch relatively infrequently, i.e. at the same rate as the environment switches. In contrast, when environment durations *T* are short, fast growing cells can still come to dominate the population when *r* is sufficiently large simply because the phenotypes of faster growing cells are more stable than those of more slowly growing cells, i.e. as previously identified in [30, 31]. In this regime, fitness is optimised by maximising switching rates.

As shown in Suppl. Figs. S5 and S6, the boundary between the long-T regime where *ϕ* = 1/*T* is optimal and the short-*T* regime where *ϕ* → ∞ is optimal, approximately corresponds to the line *T* = *r/n* when *T* ≥ 10 generations. When *T* is small, high average growth rates can only be achieved using the short-*T* strategy and require *r* ≫ *n*. When both *T* is small and GRDS is weak (small *r*), high average growth rates can not be achieved.

### General model: GRDS always increases population fitness

Our calculations so far have quantified the benefits of GRDS under the assumption that switching rates were optimised by natural selection. Next, we investigated to what extent GRDS is also beneficial when switching rates are not optimised. First, in SI Section 2.B.1 we prove mathematically that, when environment durations are not very short and switching rates obey some weak constraints, GRDS always improves the average growth rate of bet-hedging strategies. Notably, this proof applies to the general model and is thus valid for an arbitrary number of phenotypes, growth rates, environments, and regardless of whether all environments have the same duration or whether the phenotype switching rates are optimised. This proof thus strongly suggests that GRDS is generically beneficial when environment durations are not short.

Finally, to quantify the extent of the growth rate improvement when switching rates are not optimised and to explore whether the benefits of GRDS also extend to the regime of short environment durations, we numerically computed the effect of GRDS on average growth rate for many different parameter sets (Fig. 3). The parameter sets were picked as follows: we systematically varied the number of environments between 5 and 20, chose the environment switching probabilities *b_IJ_* uniformly at random, and varied the average environment duration from *T* = 1 to *T* = 40. The number of phenotypes m was chosen either equal, twice, or half the number of environments. For each environment the growth rate of the fastest growing phenotype was drawn randomly over a range, such that one unit of time on average corresponds to one doubling of the best phenotype. All other phenotypes were assigned growth rates at random, chosen uniformly over a range below the maximal growth rate in that environment. The switching rate *ϕ_ij_* for each pair of phenotypes was chosen randomly over a thousand-fold range. To model the effect of GRDS, we then add growth rate dependence to these randomly chosen switching rates, such that we get rates 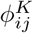 that are higher in environments where phenotype *j* grows slow, and lower where phenotype *j* grows fast (see the Methods-section and SI Sect. 3 for details). GRDS of strength *r* (plotted on the horizontal axis in Figure 3) was then implemented by making the maximal ratio between the 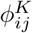 in different environments *K* equal to *r*. For *r* = 1 we thus model a population with fixed, randomly-chosen switching rates, and with *r* > 1 we model a population in which some GRDS was added.

**Figure 3.**
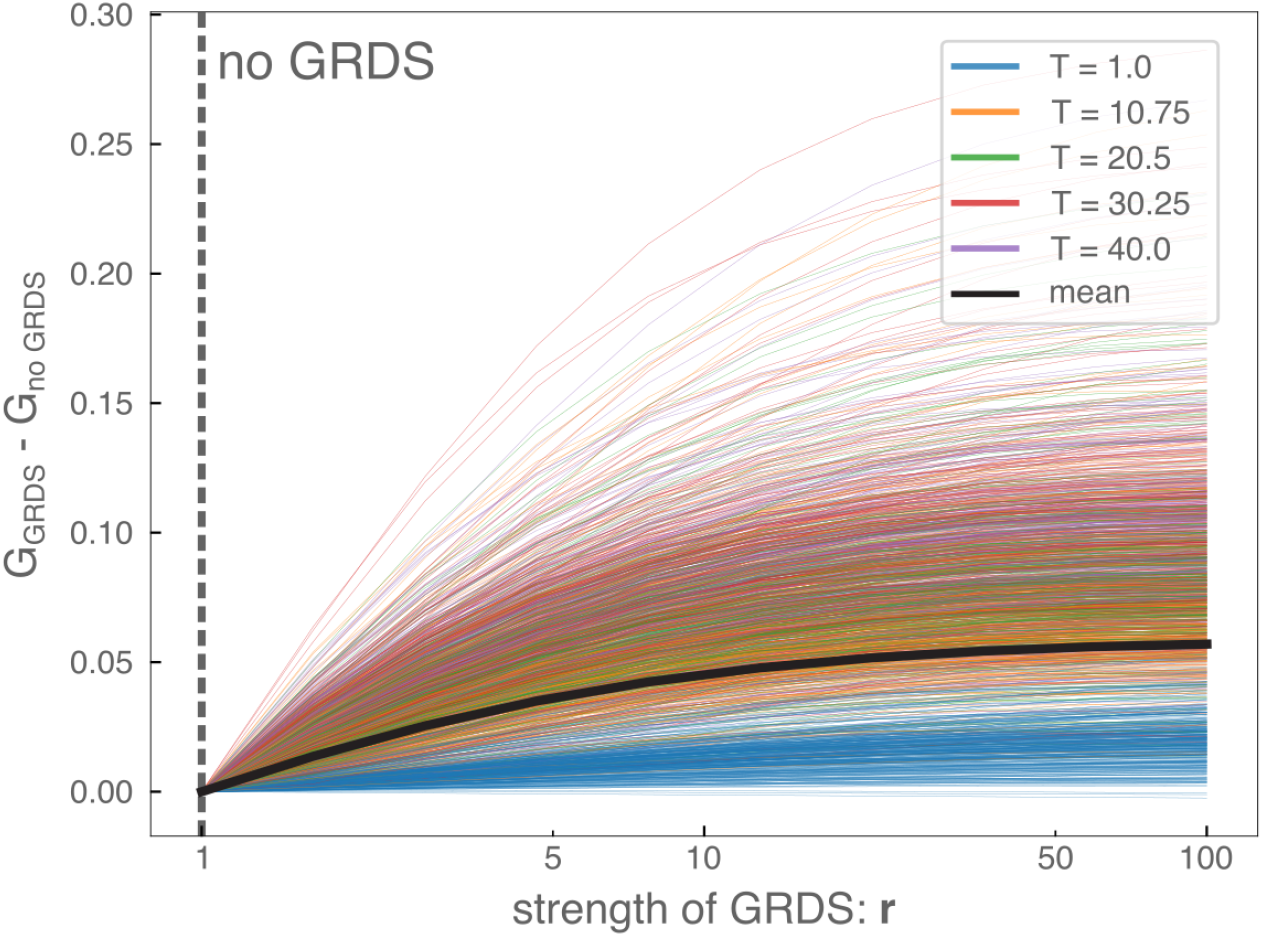
Population growth rates rise with increasing strength of GRDS *r*, even when switching rates are not optimised. Each coloured line corresponds to a different randomly-picked parameter set, and shows how the average population growth rate *G* increases when the strength of GRDS *r* is increased. Different colours correspond to different average environment durations *T*, where *T* =1 corresponds roughly to a single doubling of the fastest growing phenotype in each environment (see SI Sect. 3). The bolder black line indicates the average. The vertical axis shows the difference between the average population growth rate with a certain strength of GRDS and the average growth rate without GRDS. The rate of switching from phenotype *j* to *i* decreases with the growth rate of phenotype *j*, and the ratio of the highest and lowest switching rate from *j* to *i* in different environments is *r* (see Methods for details). The random sampling of the other parameters is described in SI Section 3.

For almost all of the parameter sets (1872 out of 1875), a small growth rate dependence immediately increases the long-term growth rate that is achieved by a randomly switching population (Fig 3). Although the size of the fitness benefit of GRDS as a function of *r* varies across the randomly chosen parameter settings and is consistently smaller when the average environment duration is short, essentially all curves show an initial steep increase followed by a slower but non-saturating increase with *r*, similar to the dependence observed for the minimal model with optimal switching rates (Equations (6) and (8)).

## Discussion

Although it has long been observed that microbial populations of isogenic cells can exhibit significant phenotypic variability, over the last two decades it has become increasingly appreciated that such phenotypic variability is pervasive and involves both continuous fluctuations in gene expression and stochastic switching between discrete phenotypic states [9–11,13,34–38]. A large body of theoretical work has shown that such phenotypic heterogeneity can be beneficial in fluctuating environments, which led to the suggestion that microbial populations may be employing ‘bet-hedging’ strategies [34, 39, 40]. However, previous theoretical work has also suggested that bet-hedging strategies can only be effective if environments are not too diverse and environment durations are relatively long [10, 18, 19]. These fairly restrictive bounds on the benefits of bet-hedging raise the question to what extent the pervasive phenotypic heterogeneity that is observed in microbial populations can be attributed to bet-hedging. Here we have shown that these bounds on the effectiveness of bet-hedging strategies disappear when we account for one additional ingredient: Growth Rate Dependent Stability (GRDS). With GRDS, phenotype switching rates decrease with growth rate and we have shown that, as the ratio between switching rates of slow- and fast-growing cells increases, the intrinsic costs of bet-hedging can be compensated and average population growth rates can approximate the theoretical maximum.

There is in fact significant evidence supporting that phenotype switching rates tend to decrease with growth rate. In a number of studies it has been observed that gene expression noise levels decrease with growth rate [22–24, 30] and metabolic heterogeneity has also been observed to increase with nutrient limitation [41–46]. Since phenotype switches are often ultimately driven by fluctuations in gene expression or metabolic state [12, 25–29, 47], phenotype switching rates will generally increase with gene expression noise levels. Notably, if we assume that a particular phenotype switch occurs under a particular rare fluctuation in gene expression, then even a small change in noise level can have a large effect on the phenotype switching rate. For example, when noise levels differ two-fold between fast and slow-growing cells, and the gene expression fluctuation that is required for a phenotype switch corresponds to 4 standard-deviations in fast growing cells, then the same fluctuation would correspond to 2 standard-deviations in slow growing cells, leading to a *e*^4/2^/*e*^22/2^ ≈ 400 fold increase in switching rate in slow growing cells.

It is currently not clear what mechanisms underlie the decrease of gene expression noise with growth rate. Analysis of genome-wide noise in *E. coli* across different growth conditions has shown that while relative noise levels of different genes are highly condition-dependent and are driven by noise propagation through the regulatory network, absolute noise levels decrease systematically with growth rate in a way that appears to affect all genes [24]. This suggests that the overall decrease in gene expression noise with growth rate results from mechanisms that affect all genes. However, this still leaves many possible mechanisms including fluctuations in chromosome copy numbers across the cell cycle, fluctuations in transcription initiation rates due to variations in RNA polymerase concentration, fluctuations in transcription elongation rates due to variation in nucleotide concentrations, fluctuations in translation initiation rates due to variation in ribosome concentration, fluctuations in translation elongation rates due to variation in charged tRNA concentrations, fluctuations in dilution rate due to variation in growth rate, intrinsic Poissonian fluctuations in all steps of the gene expression process, unequal division of proteins at cell division, and so on. Although a number of models has been proposed that show how some of these sources of noise may explain a decrease in noise with growth rate, e.g. [22,31,48,49], these models make many simplifying assumptions and only consider some of the mechanisms listed above. As it is currently unknown which mechanisms are most important for determining expression noise levels in realistic settings, it is thus not yet clear which mechanisms drive the observed decrease in noise with growth rate.

We hypothesise that one important contributor to the decrease of noise with growth rate is that the growth rate sets the dilution rate of most intracellular molecules and thus also the rate at which intracellular fluctuations are diluted. Although both the frequency and amplitude of some intracellular noise may naturally increase with growth rate, thereby compensating for the increased dilution rate, this may not apply to all sources of noise, such as fluctuations in extracellular levels of metabolites and stressors. Indeed, in a study of regulatory circuits with positive feedback in *E. coli*, we have recently shown that, because signalling molecules are diluted more quickly at higher growth rates, the sensitivity of these regulatory circuits to external signals generally decreases with growth rate [50], supporting that faster dilution may dampen fluctuations in the internal states of cells. Of course, since GRDS generally increases long-term average growth rates, the decrease of fluctuations with growth rate may even be an adaptation that has evolved through natural selection.

Finally, we have so far implicitly assumed that sensing/regulation and bet-hedging are mutually exclusive strategies, but these strategies can of course act in parallel and may in fact be deeply entangled. By comparing native and synthetic promoters, we have previously shown that natural selection has acted to increase the noise levels of native *E. coli* promoters [51]. Moreover, expression noise in *E. coli* results to a substantial extent from noise propagating through the regulatory network so that noise levels are highly condition-dependent, with noise in more-regulated promoters being both higher on average and more variable [24]. These observations indicate that gene regulation and expression noise are intimately coupled and that the fluctuations in gene expression that we call ‘noise’ at least to some extent result from fluctuations in environmental conditions that propagate through the gene regulatory network. This suggests a strategy in which sensing, regulation, and bet-hedging are all acting in concert, with sensing and regulation being used to constrain the subspace of phenotypes that is explored by stochastic phenotype switching.

## Methods

The general setup of our model has been introduced in the ‘Model Setup’-part of the Results-section (see also SI 2.A.1). We have studied this model through several methods: by simplifying it to a toy example and a minimal model, by mathematically studying a limit where environment changes are relatively infrequent and switching rates are low, and by simulating the general model for many random samples of the model parameters. We will here briefly describe each of these methods. All mentioned `Python`- and `Mathematica`-scripts have been made publicly available at https://github.com/dhdegroot/GRDS-code-repository.

### Toy example

For Figure 1 we simulated the general model using a `Python`-script with only *n* + 1 = 3 phenotypes in a sequence of two environments with a duration of *T* = 10. The growth rate of the optimal phenotype was set to *μ*_1_ = 1, while the other phenotypes did not grow: *μ*_0_ = 0; the strength of GRDS was *r* = 10. For bet-hedging cells, we allow for only one global switching rate: 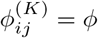. With GRDS, this switching rate becomes *rϕ* from non-growing phenotypes. In both cases *ϕ* was numerically optimised to maximise average growth rate.

### Minimal model

All results related to the minimal model were obtained with `Mathematica` and an analytical expression was obtained for the average growth rate *G* as a function of the parameters (*n, T, r, ϕ*). Optimisation of the switching rate *ϕ* to maximise the average growth rate *G* was done numerically.

### Mathematical proofs

In Supporting Information Section 2.B.1 we prove that GRDS can always increase the average population growth rate *G*; in Section 2.B.2 we approximate the fitness benefit as a function of the strength of the growth rate dependence *r*. These proofs are only possible if we apply the same approximation as proposed in [18], which entail:

1. The duration of environments is long enough compared to division times and switching times, such that the phenotype distribution in the population has relaxed to a stationary distribution before the next environment switch.
2. The switching rates are small compared to the differences between the fastest growth rate and other growth rates in an environment, such that the stationary phenotype distribution can be well-approximated by determining the dominant eigenvector of the time-evolution matrix with perturbation theory.

### Numerical simulations

The numerical calculations for the general model were done in `Python`. As detailed in SI Sect. 3, we randomly pick a number of environments *m*, a number of phenotypes *n*, an average environment duration *τ*, a growth rate for each phenotype in each environment 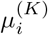, and parameters *b_IJ_* that determine the random sequence of environment-types. Then we randomly choose a switching rate *ϕ_ij_* for each pair of phenotypes *i, j*, and implement GRDS of strength *r* as follows. We start by taking the range *R_ϕ_*(*r*) = [log(*ϕ_ij_*) – 0.5log(*r*),log(*ϕ_ij_*) + 0.5log(*r*)]. Then we determine the range of growth rates that cells of phenotype *j* can achieve in the different environments: 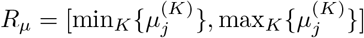, and we let *t^r^* be the linear map from *R_μ_* to *R_ϕ_*(*r*) that maps the minimal growth rate to the maximal switching rate and vice versa. Now the switching rate from phenotype *j* to *i* in environment *I* with GRDS of strength *r* is determined by 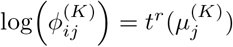. Based on the growth rate in the current environment, the switching rates between two phenotypes are thus linearly interpolated in log-scale between an upper bound and a lower bound, where the factor difference between the upper and lower bound is *r*. In addition, to allow an unbiased comparison between different strenghts of GRDS, we rescale these switching rates such that the average switching rate between two phenotypes 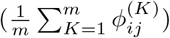 is equal to the initially drawn switching rate *ϕ_ij_* for all strengths of GRDS. For the results presented in Figure 3, we did these simulations for 10 different values of *r*, where *r* = 1 corresponds to the case with no GRDS.

Given this complete set of parameters, we compute the average population growth rate by simulating a sequence of environments with a total duration that exceeded *n*^2^*τ* to ensure that we sufficiently sampled the possible switches between different environments.

## Acknowledgements

DHdG and AJT thank Gašper Tkačik for his useful input. This work was supported by NWA grant 1228.191.283 from the Dutch Research Council and SNF grant 310030_184937 from the Swiss National Science Foundation.

## Contributions

D.H.dG., A.J.T., and E.vN. designed the research. D.H.dG. and A.J.T. performed the research. F.J.B. and E.vN. supervised the research. D.H.dG., A.J.T., F.J.B, and E.vN. wrote the paper.

## Supplementary Information

The Supporting Information Text is organised as follows. In Section 1 we describe the setup and analysis of our minimal model; this section is complemented by the Mathematica notebook named `GRDS_minimal_model_analysis.nb`. In Section 2, we derive an approximation of the average population growth rate under the assumption that the environment durations are relatively long, and the switching rates relatively small. We prove that GRDS always increases the average population growth rate as long as this approximation holds. Finally, in Section 3 we describe how we performed numerical simulations that test if the results obtained in Sections 1 and 2 hold for the most general version of the model.

### 1 Analytical derivations minimal model

#### 1.A Model set-up

In this first part of the SI, we investigate a minimal model of Growth Rate Dependent Stability (GRDS). We start by assuming that there are only two growth rates with *μ*_1_ > *μ*_0_, and that in each environment there are *n* slow-growing phenotypes for only 1 fast-growing phenotype. Cells can stochastically switch between phenotypes, and we will (for now) assume that the switching rate between all phenotypic states is equal: *h/n*, which is chosen such that the total switching rate away from each phenotype is equal to *h*. Growth rate dependence of the switching rates will be introduced later. In one environment, we thus get the set of differential equations:

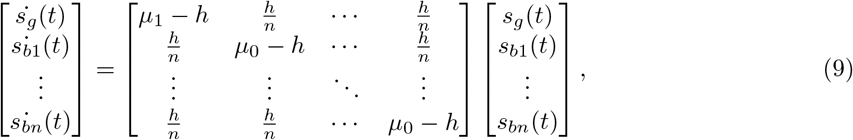

#### 1.B Rewriting the model

##### 1.B.1 Summing the cells in slow growth phenotypes

We will show that we can safely approximate this system as a two-state system, where we describe only the sum of all the cells that are in a slow-growth phenotype instead of all phenotypes separately. From (9), we can write down the differential equation for the sum of all cells in the slow growth states:

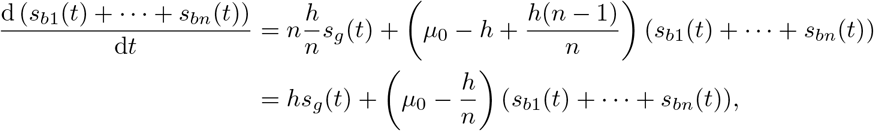

and for the cells in the fast growth states we get:

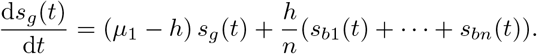

So, if we define *s_b_*(*t*) as the number of cells that are in one of the bad states *s_b_*(*t*) = *s*_*b*1_(*t*) + ⋯ + *s_bn_*(*t*) then we can write

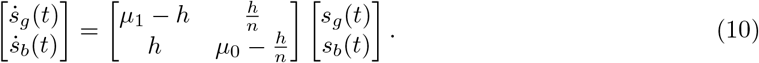

Note the intuition here: the switching rate from all states is *h*, but the probability that a cell that switches from a bad phenotype ends up in a good phenotype is 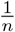 since there are *n* – 1 different bad phenotypes and only one good phenotype. The net switching rate from the bad to the good phenotype is therefore 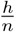.

##### 1.B.2 Shifting the slow growth rate to 0

To make our model as minimal as possible, we can rescale the system to get rid of some parameters. First, let us check what the effects are of shifting both growth rates by a constant, i.e. we use *μ*_1_ → *μ*_1_ – *δμ* and *μ*_0_ → *μ*_0_ – *δμ*. Let us assume that we already have a solution 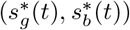 that solves the system in (10). We can use this to generate a solution for the shifted ODEs

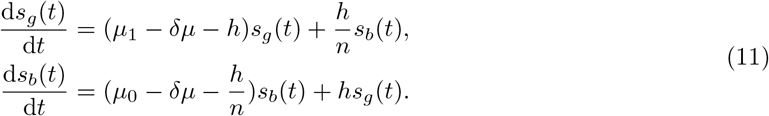

To see this, take 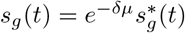 and 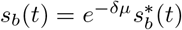. If we insert this in the differential equation for *s_g_*(*t*), we get:

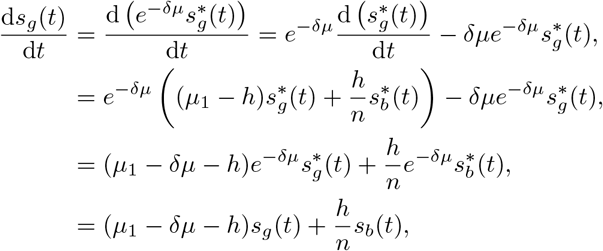

and similar for the differential equation of *s_b_*(*t*). This shows that the solutions are only rescaled by an overall factor when we shift the growth rates by an equal amount. Intuitively, the reported growth rates in the new system are just lower by *δμ*, but the relative growth in the different phenotypes is the same.

We can now choose *δμ* = *μ*_0_. This implies that we can always look at systems where the slow growth rate is zero, without loss of generality. We get

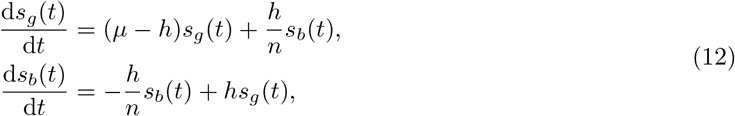

where we have used *μ* = *μ*_1_ – *μ*_0_.

##### 1.B.3 Scaling the fast growth rate to 1

We now still have a degree of freedom in picking our time units. We can use the fast growth rate, and take *τ* = *μt*, which means that one time unit (*τ* = 1) now corresponds to 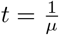, which is the time that it takes a fast growing cell to grow to a factor *e*. The benefit of this rescaling is apparent when we insert it in the differential equations, for example

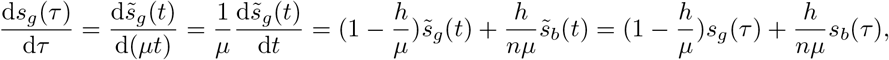

where 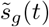 is defined such that 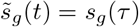. We can now define 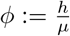 as the rate of switching relative to the growth rate of the fast growing phenotype. One can also look at it from another perspective by writing 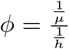, which can be interpreted loosely as the ratio of the cell cycle time with the time it takes to switch, or: how often does a cell switch before it doubles. When we again write *t* for our rescaled time instead of *τ*, we get our minimal differential equations:

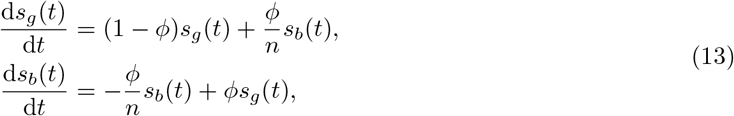

#### 1.C Solving the differential equations

##### 1.C.1 The general analytical solution to the differential equations

The system in (13) can be solved analytically by solving the quadratic equations to find the eigenvalues and -vectors. The solution can be found in the Mathematica notebook: `GRDS_minimal_model_analysis.nb`.

Since we are eventually only interested in the growth rate, we have some freedom in choosing the initial number of cells. We choose the total number of cells to be 1 at timepoint 0: *s*_tot_(0) = *s_g_*(0) + *s_b_*(0) = 1. We then know that a fraction of the population is in the good phenotype, *s_g_*(0) = *p*_0_, and the rest is in one of the bad phenotypes *s_b_*(0) = 1 – *p*_0_. Given these initial conditions, we can find out the state of the population after one environment duration: *T* (note that *T* is now in units that give the times that a fast growing cell could have e-folded itself). We get expresssions for *s_g_*(*T, ϕ, n, p*_0_) and *s_b_*(*T, ϕ, n, p*_0_). At the end of the period, we can again calculate the fraction of adapted cells:

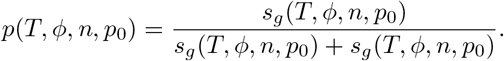

The analytic expression for this fraction is fairly cumbersome (see Mathematica-file `GRDS_minimal_model_analysis.nb`), but if we gather a recurring combination of terms in a separate variable, *D*, we can write it down as:

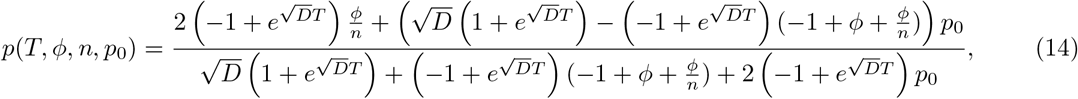

where

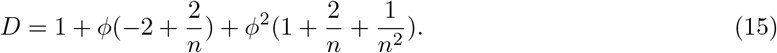

We find that the large-*t*-limit is independent of *p*_0_, as expected:

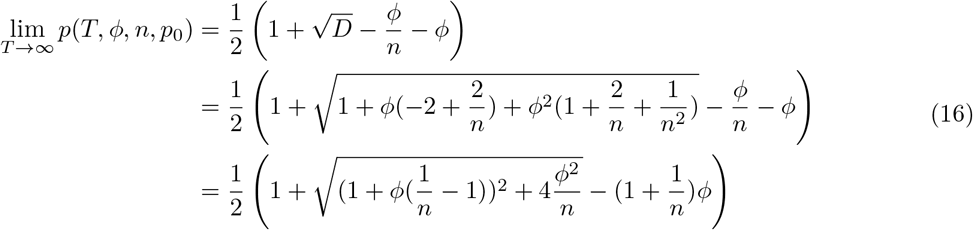

##### 1.C.2 Introducing periodic initial conditions

When the environment switches, we assume that all cells in a good state will automatically switch to a bad state. However, the cells that were in a bad state might coincidentally be pre-adapted for the new environment. We will now make the assumption that the non-adapted cells are distributed equally over the non-adapted phenotypes at the end of an environment, and we will discuss below why this is reasonable for a very large parameter regime. Under this assumption, since there are *n* phenotypes, a cell in a bad phenotype has a probability of 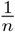 to be pre-adapted upon an environment switch, and the initial fraction of cells in a good phenotype will thus be 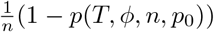. This can be seen as a recursive mapping: 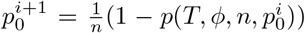. We prove below that this initial fraction of adapted cells will converge to a fixed point 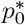 upon repetition of this mapping.

###### Lemma 1.

*Let the time evolution within one environment of the number of adapted, s_g_*(*t*), *and nonadapted, s_b_*(*t*), *cells be given by* (13). *The fraction of adapted cells in the i-th environment is given by*

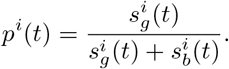

*Say that, after a fixed environment duration T, the environment switches and the new fraction of adapted cells is given by* 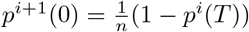. *Then, the initial fraction of adapted cells will converge when the number of past environments becomes large*: 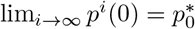.

*Proof.* The time evolution of the number of cells can be captured as a linear system of differential equations:

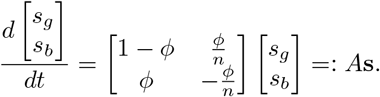

If the environment stays for a certain fixed time *T*, we can solve this equation as **s**(*T*) = **s**(0)*e^AT^*, where *e^AT^* is again a matrix. Upon an environment switch, the redistribution of the cells over adapted and non-adapted phenotypes can also be written as a matrix multiplication:

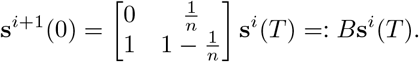

Together this shows that for a fixed environment duration *T* we can write down a linear map from the cell numbers at the start of the first environment, to the cell numbers at the start of the second environment, and so on:

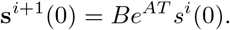

Now let us consider the matrix *Be^AT^*. One can check that the assumptions for the Perron-Frobenius theorem hold for this matrix, i.e., there is some λ such that *Be^AT^* + λ*I* is a positive-valued matrix that is primitive (see Section 2.A.2 for details where we check this for a very similar matrix). The Perron-Frobenius theorem tells us that *Be^AT^* has a real-valued largest eigenvalue and that there is a nonzero difference to all other eigenvalues. This means that we can use the diagonalisation of the matrix to write it as *Be^AT^* = *MΛM*^-1^, where *M* is the matrix with the eigenvectors of *Be^AT^* and Λ is the diagonal matrix with the eigenvalues. If we now want to know the initial numbers of cells after *N* iterations of the environment we can write:

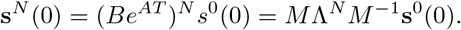

Because the dominant eigenvalue is larger than all other eigenvalues, only the corresponding (dominant) eigenvector will determine the *s^N^*(0) for large enough *N*. Since the initial fraction of good cells is completely determined by this *s^N^*(0) by 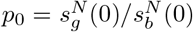 we thus know that this will stabilise to a fixed number after enough environment iterations.

We thus know that eventually there is a fixed initial fraction of adapted cells that satisfies the following equation (as illustrated in Figure S1):

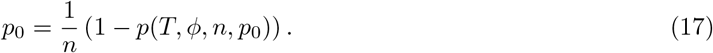

**Figure S1.**
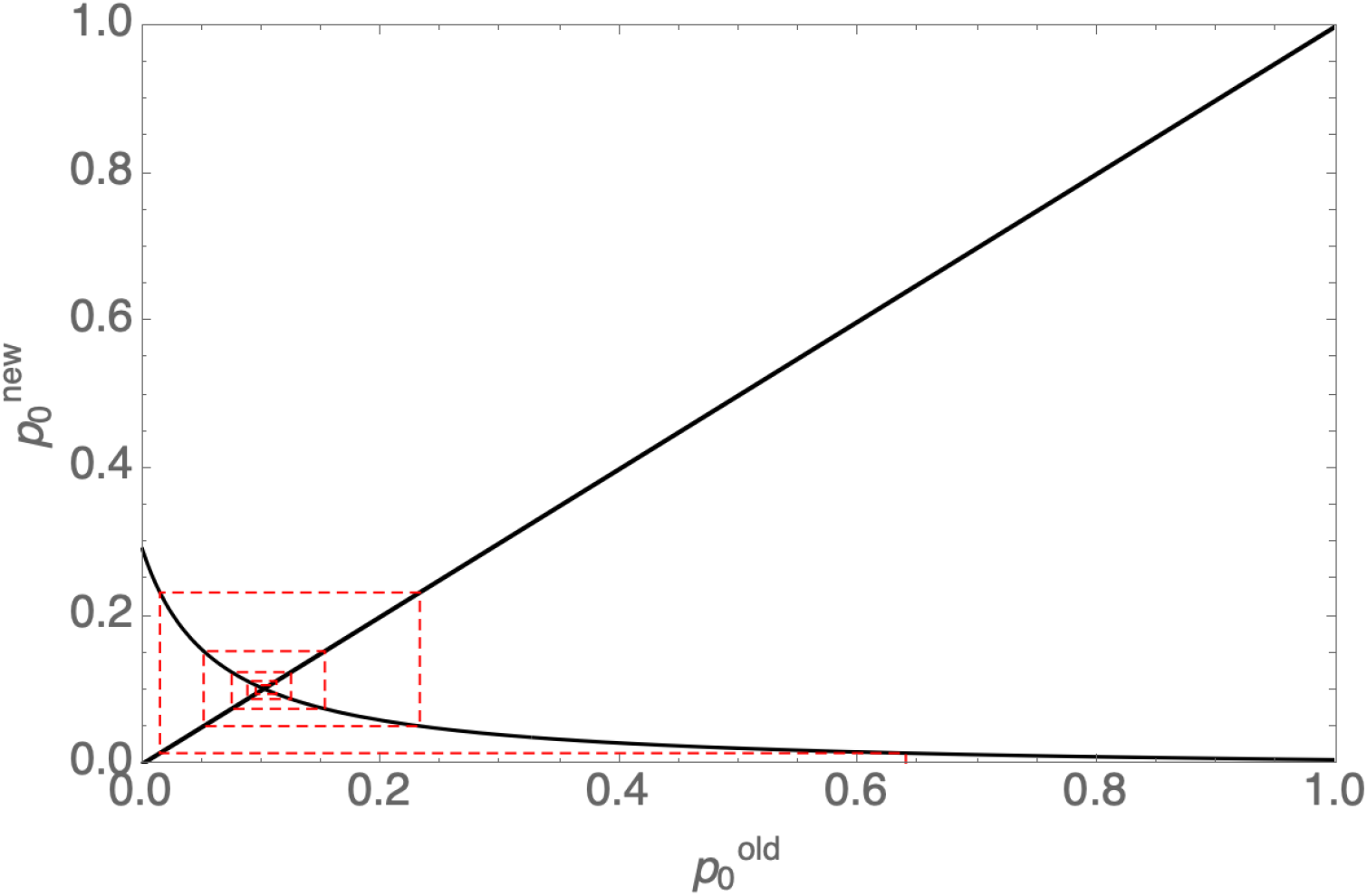
The initial fraction of adapted cells converges to a fixed point after several iterations of environment changes. The new initial fraction of adapted cells depends on the eventual fraction of adapted cells through: 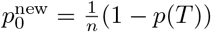. Since, *p*(*T*) can be expressed in terms of 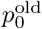, the relation between 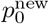 and 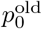 can be plotted. The red, dashed line, shows how the initial fraction converges to a fixed point after several iterations. This figure was made for parameters: *ϕ* = 0.02, *n* = 3, *T* = 3, 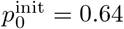, using the Mathematica-notebook `GRDS_minimal_model_analysis.nb` which we have made publicly available.

The solution of (17) can be found in the Mathematica-notebook `GRDS_minimal_model_analysis.nb`. Since we already had a solution for the total number of cells *s*_tot_ (*T, ϕ, n, p*_0_) = *s_g_*(*T, ϕ, n, p*_0_)+ *s_b_*(*T, ϕ, n, p*_0_), we can insert the solution for 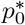 to get the growth dynamics when this fixed point is reached. We can use this to calculate the average growth rate:

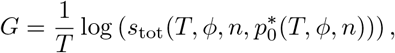

which is only a function of *T, ϕ* and *n*. This provides an analytical expression for the long term average growth rate as a function of all parameters in the model. We have not found an analytical expression for the optimal switching rate *ϕ*^opt^, but it is easy to find it numerically for a given set of parameters.

###### The approximation that non-adapted cells are equally distributed over non-adapted phenotypes

Above, we have used the approximation that at the end of an environment the non-adapted cells are distributed equally over the non-adapted phenotypes. This is a very good approximation as long as the environment durations are not very short, or the switching rates very low. Indeed, although the cells may be non-equally distributed over the non-adapted phenotypes at the start of a new environment (because one of these non-adapted phenotypes was the adapted phenotype in the previous environment), they are re-distributed via two different mechanisms. First, non-adapted cells switch between each other with the same rate *ϕ* (which will become *rϕ* once we introduce GRDS). This switching will re-distribute the cells equally over the non-adapted phenotypes. As long as most non-adapted cells have switched at least once during the current environment, cells will have redistributed randomly over the phenotypes and this occurs when *rϕT* is on the order of 1 or larger, i.e when *rϕ* is not small compared to the environment switching rate 1/*T*. Alternative, the non-adapted phenotypes may also be equilibrated by switching of adapted cells to non-adapted phenotypes. Note that the cells in the adapted phenotype will start growing and their offspring will switch to all non-adapted phenotypes at the same rate *ϕ/n*. When the environment duration is large enough, the cells in the good phenotype have produced enough offspring that switched to non-adapted phenotypes to equilibrate the cell numbers in those phenotypes. To make this specific, at the start of an environment there will be one non-adapted phenotype with many more cells than the others because this phenotype was optimal in the previous environment. Generally, it will have *N*(1 – *ϕ*) cells, while the other phenotypes each contain *Nϕ/n* cells. One of the other phenotypes is adapted to the current environment, and these cells will grow to approximately 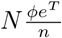 after a time *T*. By that time 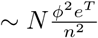 of these cells will have switched to each of the other phenotypes. If 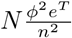 is large compared to *N*(1 – *ϕ*), these switched cells will have equalised the initial difference in cell number in the non-adapted phenotypes. From this we see that whenever *T* is larger than 2 log(*n/ϕ*), the non-adapted cells are also guaranteed to have equilibrated.

Finally, note that separating these two mechanisms of re-distributing the non-adapted cells leads to quite pessimistic bounds on when the approximation is reasonable. In reality, these two mechanisms will work in concert, so that the approximation is reasonable for a considerably larger range of parameters. As confirmed by the contour plots of Figures S5 and S6, there is indeed only a small region for small *r*, short *T*, and large *n*, where the optimal switching rates behave qualitatively different. In this regime the approximation breaks down and any results near this regime are unreliable. However, note that in this regime bet hedging strategies are guaranteed to perform poorly in any case.

##### 1.C.3 Summarising the model of conventional bet hedging

We have here reduced a model of conventional bet hedging to its simplest form, (13), after which we introduced periodic initial conditions. For any choice of the parameters *n* and *T*, and a given switching rate *ϕ* we have an analytical expression for the long-term average population growth rate *G*. This allows us to numerically optimise the switching rate to maximise *G*.

#### 1.D Introducing growth rate dependence

We will now introduce growth rate dependent stability (GRDS) in the model. Let’s recall the original system of equations here:

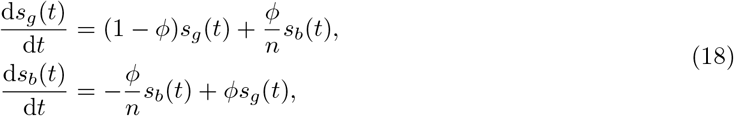

We now introduce Growth Rate Dependent Stability by allowing for two different switching rates: *ϕ* determines the switching away from adapted phenotypes, and *ϕ*_0_ = *ϕr* denotes the switching rate away from non-adapted phenotypes. The parameter *r* > 1 thus captures the strength of GRDS by giving the factor difference between fast and slow switching rates. This gives the following system

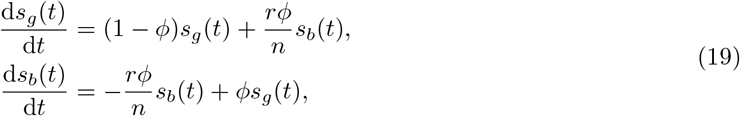

The new system ((19)) is of the same form as the original system ((18)) if we use: 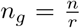, such that 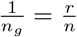. This new parameter may, just like the original 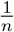, be interpreted as a parameter that captures the decrease in switching rate from a bad phenotype to the good phenotype. This gives

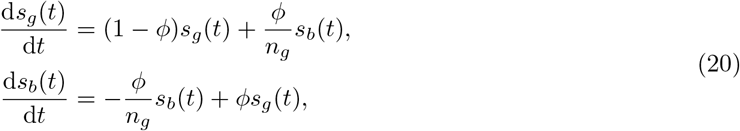

Since this model is essentially the same as before, we can re-use (14), (15), and (16). The limiting fraction for large *T* for example becomes

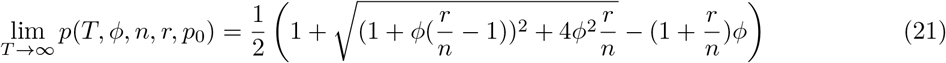

Figure S2 shows that the growth rate dependent stability already has a large effect on this limiting fraction of cells in the good phenotype.

This new GRDS model behaves essentially different from the model of conventional bet hedging when we look at the periodic initial conditions. These initial conditions are still given by the fixed point equation: 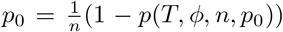, and Theorem 1 is easily extended to show that this fixed point is again globally stable. However, note that the equation is still dependent on 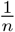, and not on the new parameter 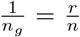. Thus, Growth Rate Dependent Stability can overcome the trade-off between fast adaptation (high switching rate needed from non-adapted phenotypes) and a fast stationary growth rate (low switching rate needed from adapted phenotypes), but it does not avoid that a high fraction of adapted cells at the end of one environment leads to a small adapted fraction at the start of the next environment.

In the following, we will study the effect of GRDS by varying the parameter *r*, while we optimise the switching rate *ϕ* for every case.

##### 1.D.1 The dependence of *G* on the model parameters

In Figures S3 and S4 we plot how the average population growth rate depends on the complexity of the environment (captured by the number of bad phenotypes *n*), and on the duration of the environment. On the horizontal axis we vary how much Growth Rate Dependent Stability the system has: from no GRDS at *r* = 1 until a thousand-fold difference between fast and slow switching rates (*r* = 1000).

**Figure S2.**
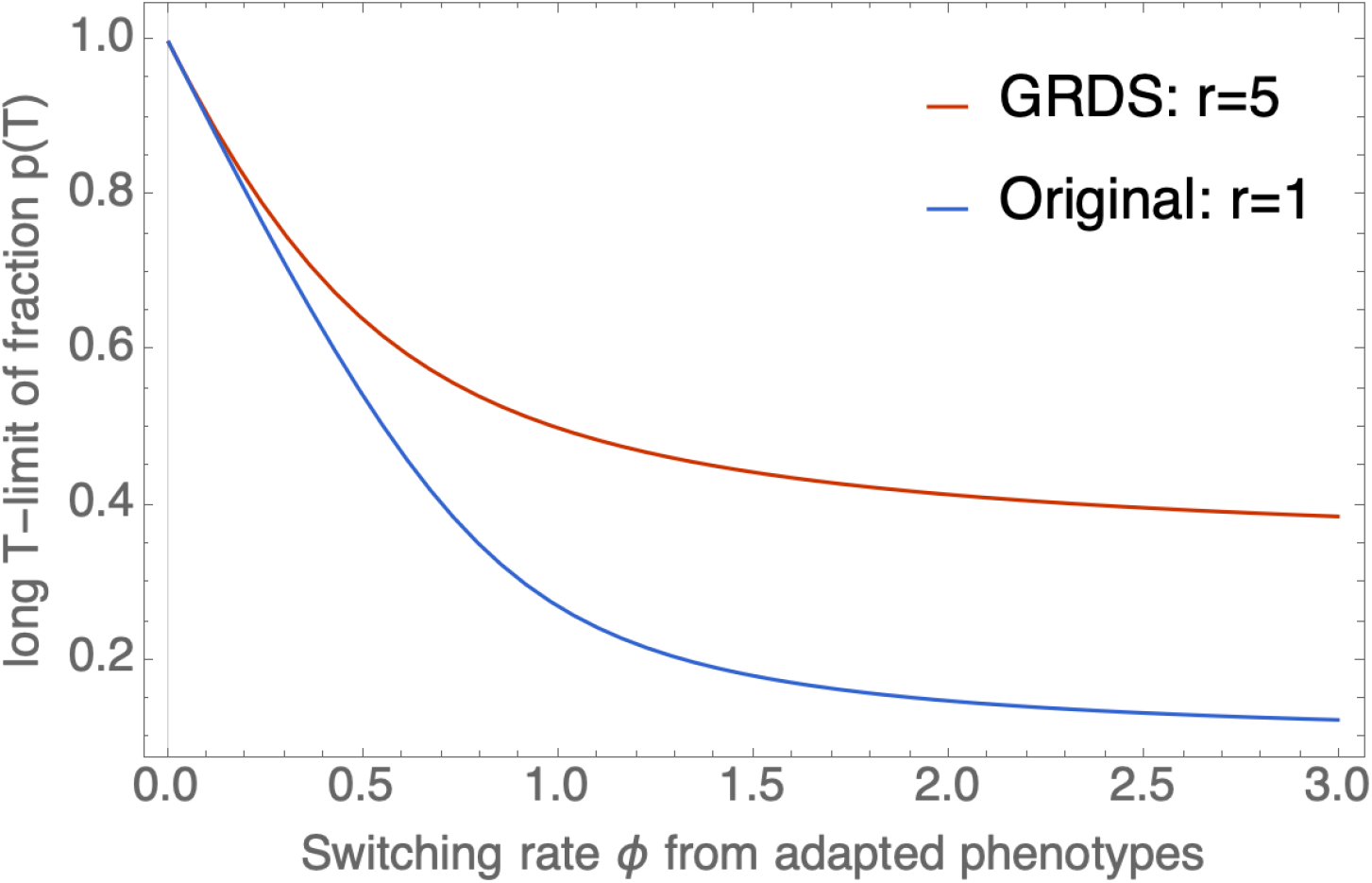
At the same switching rate away from adapted phenotypes, Growth Rate Dependent Stability keeps more cells in the adapted state. Shown is the dependence of the limiting fraction of adapted cells (for very long environment duration *T*) on the switching rate. The blue line shows conventional bet hedging; the red line shows bet hedging with GRDS of strength *r* = 5. The number of bad phenotypes was chosen to be *n* = 10. This figure was made with the Mathematica notebook `GRDS_minimal_model_analysis.nb`.

**Figure S3.**
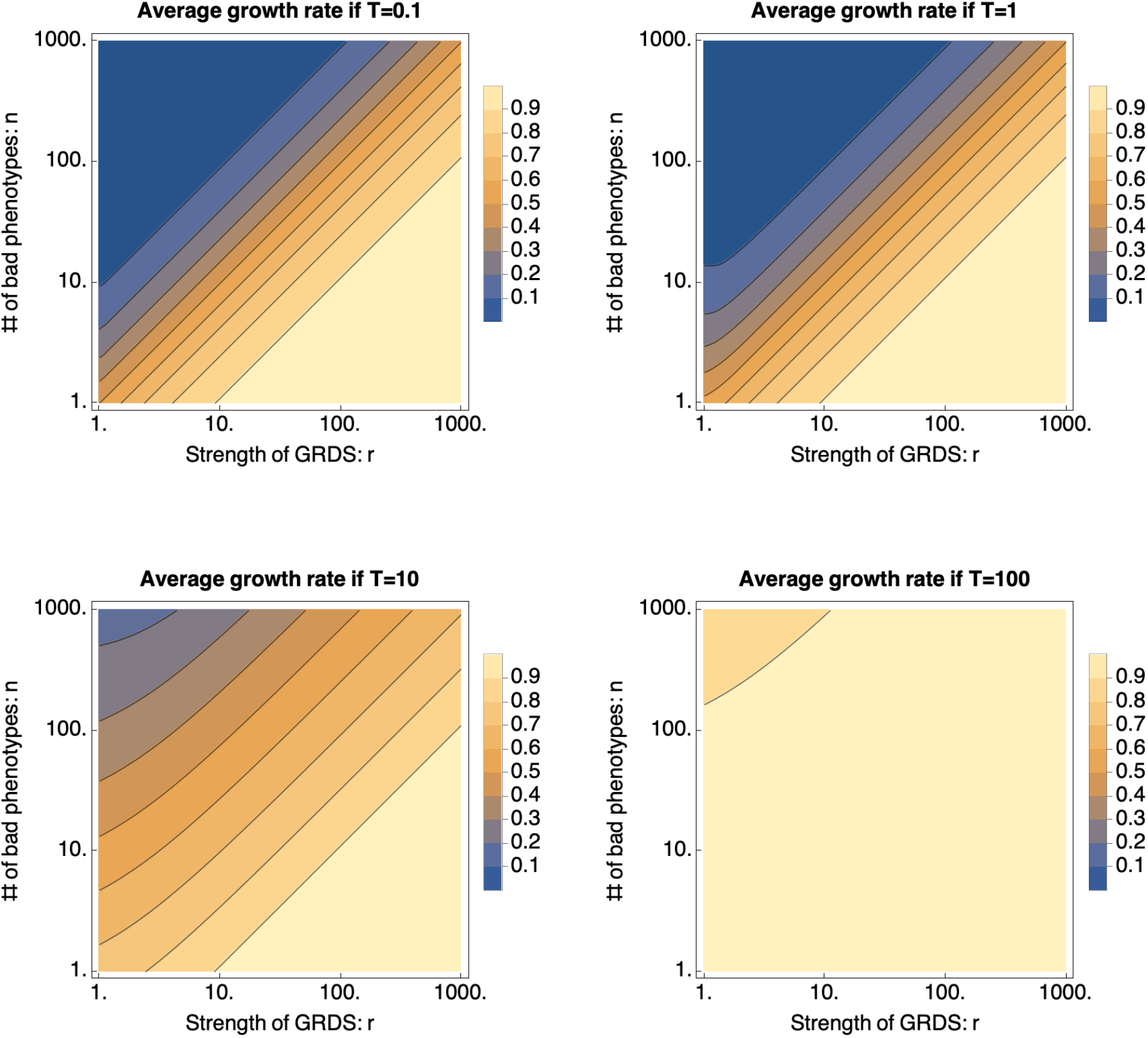
Average population growth rate *G* for a bet-hedging population that experiences GRDS. On the horizontal axes, we change the strength of GRDS (i.e. the fold-change *r* between switching rates of fast and slow growing cells). On the vertical axes, we change the number of bad phenotypes *n*. Each panel corresponds to a different environment duration *T*. The figure was made with the Mathematica notebook `GRDS_minimal_model_analysis.nb`.

**Figure S4.**
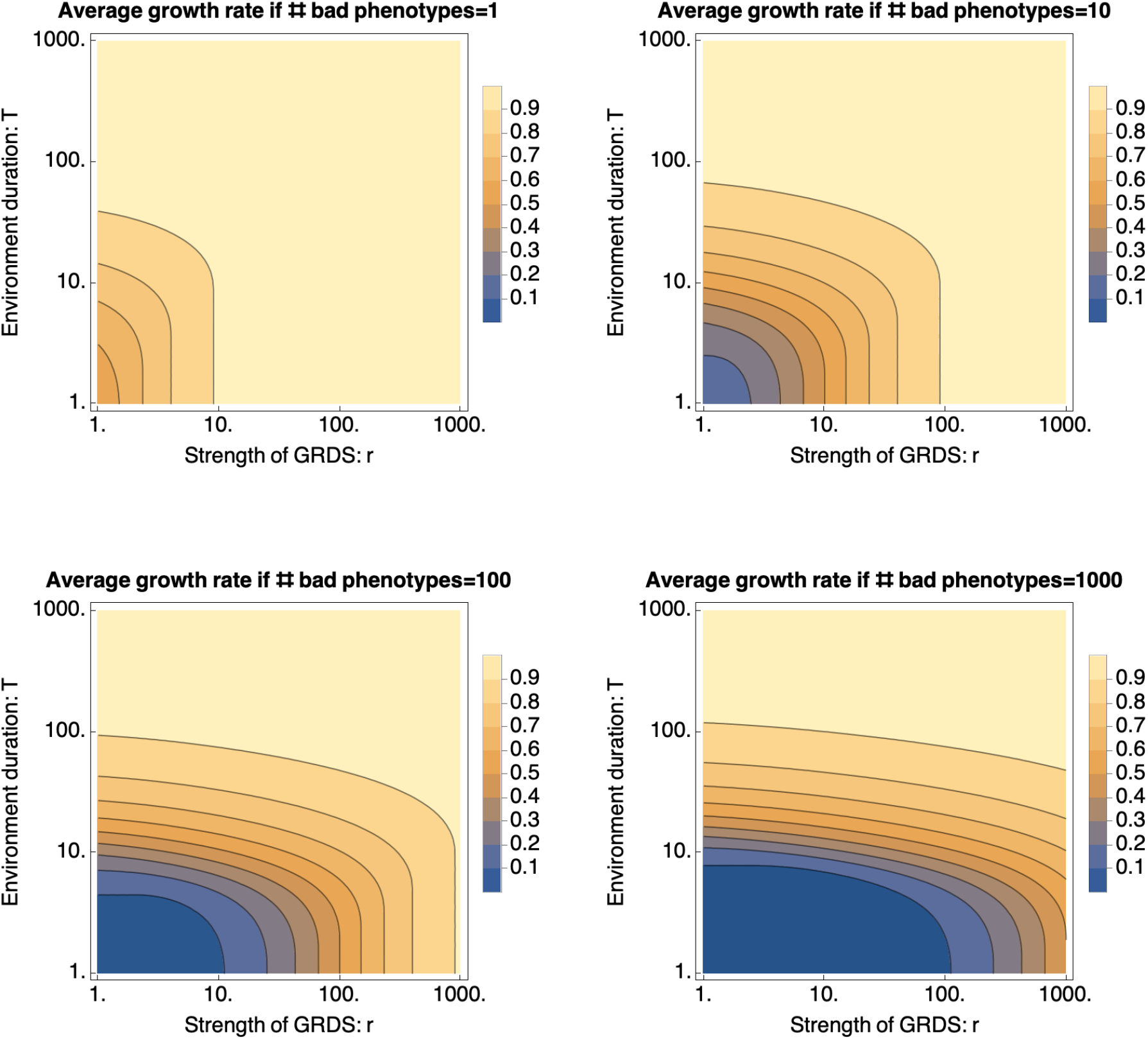
Average population growth rate for a bet-hedging population that experiences GRDS. On the horizontal axes, we change the strength of GRDS (i.e. the fold-change *r* between switching rates of fast and slow growing cells). On the vertical axes, we change the environment duration *T*. Each panel corresponds to a different numbers of non-adapted phenotypes *n*. This figure was made with the Mathematica notebook `GRDS_minimal_model_analysis.nb`.

##### 1.D.2 Model behaviour for very fast switching: large *ϕ*

For very fast switching we can get an analytical result for the average growth rate *G*. The different switching rates will determine solely what the fraction of cells in the good phenotype will be. Therefore, *p*(*t, ϕ, n*) will become 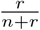 extremely fast, and the growth rate difference between cells in good and bad phenotypes will not change this fraction (because growth is slow relative to switching). Therefore, at (almost) any timepoint, a fraction of 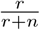 of the cells will grow with rate 1, while the rest does not grow. Therefore,

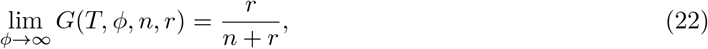

which reduces to 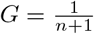 for bet-hedging without GRDS. Note that growth rate thus increases monotonically with *r* and limits to *G* = 1 for *r* → ∞.

#### 1.E Optimal switching rates

For the computation of the average population growth rates shown in Figures S3 and S4, we needed to optimise the switching rate *ϕ*. In Figures S6 and S5 we show the optimal switching rates for different parameter settings.

These contourplots indicate that, for large enough *T* (approximately *T* ≥ 10), there is an abrupt change in the optimal switching behaviour. We make this clear by plotting a red line for *T* = *r/n*. When *T* is larger than *r/n*, the optimal switching rate seems to be well-approximated by *ϕ*^opt^ ≈ 1/*T*. We check this by plotting the dependence of the average population growth rate *G* for different parameter settings in Figure S7. Indeed, in Section 2 we will prove mathematically that for long enough environment durations and relatively low switching rates, the optimal switching rate is indeed by 1/*T*. There, we also find that the average long-term growth rate is well-approximated by

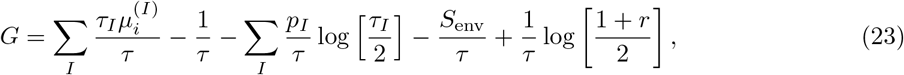

which in the case of our minimal model reduces to

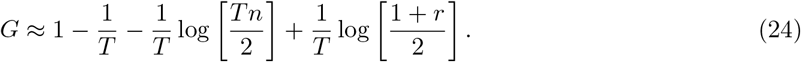

In Figure S8 we check that this expression for the growth rate is indeed a good fit as long as *r* < *Tn*, where the actual growth rate starts saturating towards the theoretical maximum of 1. In Section 2 it will become clear why the approximation breaks down when *r* becomes too large, because the switching rate from the non-adapted phenotypes, *rϕ*, becomes too large for the necessary assumptions to hold.

In the parameter regime where *T* < *r/n* we get that *ϕ*^opt^ → ∞, so that the growth rate is described by 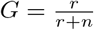 as pointed out in subsection 1.D.2.

**Figure S5.**
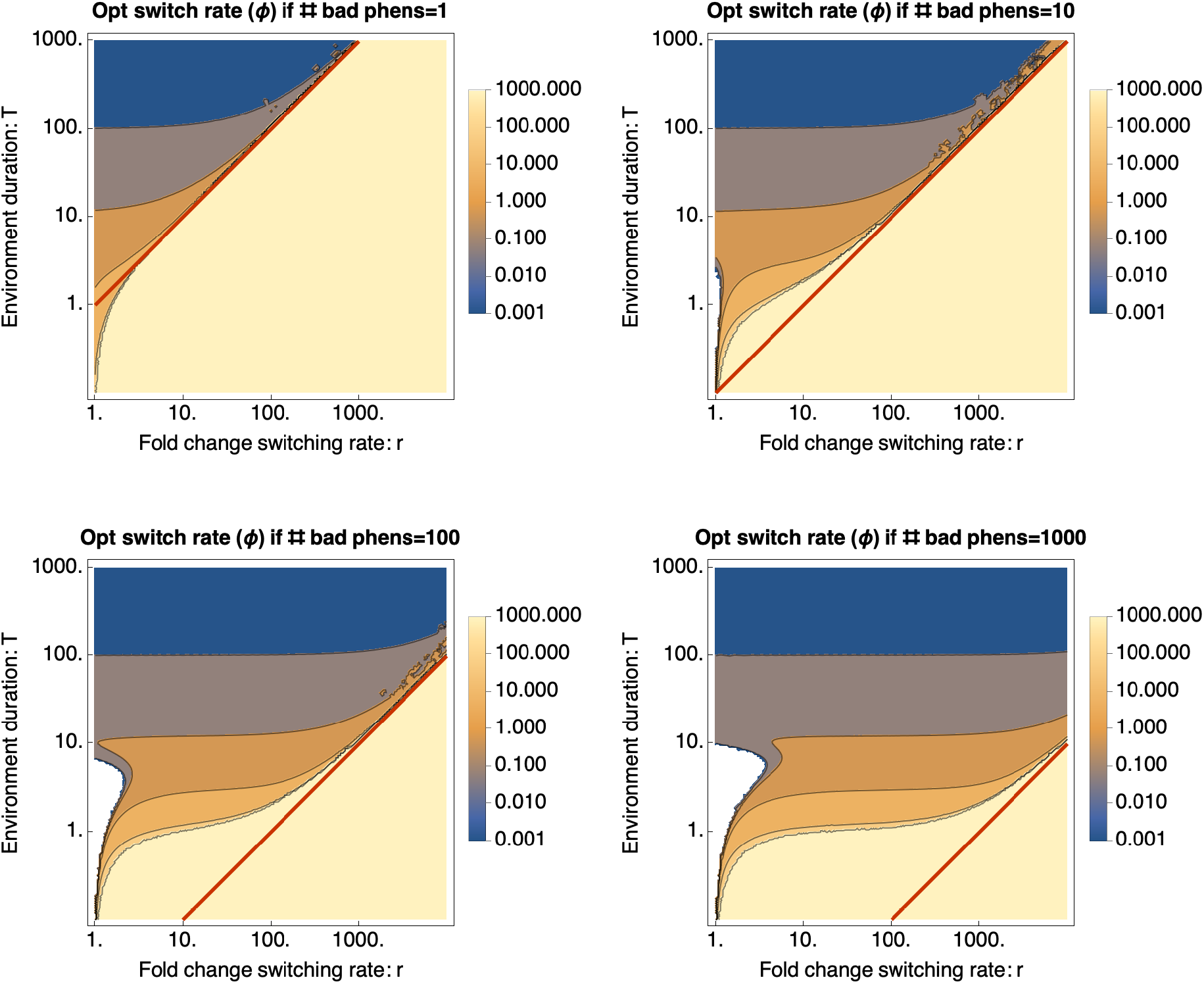
Optimal switching rate *ϕ* for a bet-hedging population that experiences GRDS. On the horizontal axes, we change the strength of GRDS, i.e. the ratio *r* of switching rates of slow and fast growing phenotypes. On the vertical axes, we change the environment duration *T*. Each panel corresponds to a different number of bad phenotypes *n*. The red line shows *T* = *r/n* which, for large enough *T*, divides the parameter regimes where *ϕ* ≈ 1/*T* (for *T* > *r/n*) and where *ϕ* → ∞ (for *T* < *r/n*). This figure was made with the Mathematica notebook `GRDS_minimal_model_analysis.nb`.

**Figure S6.**
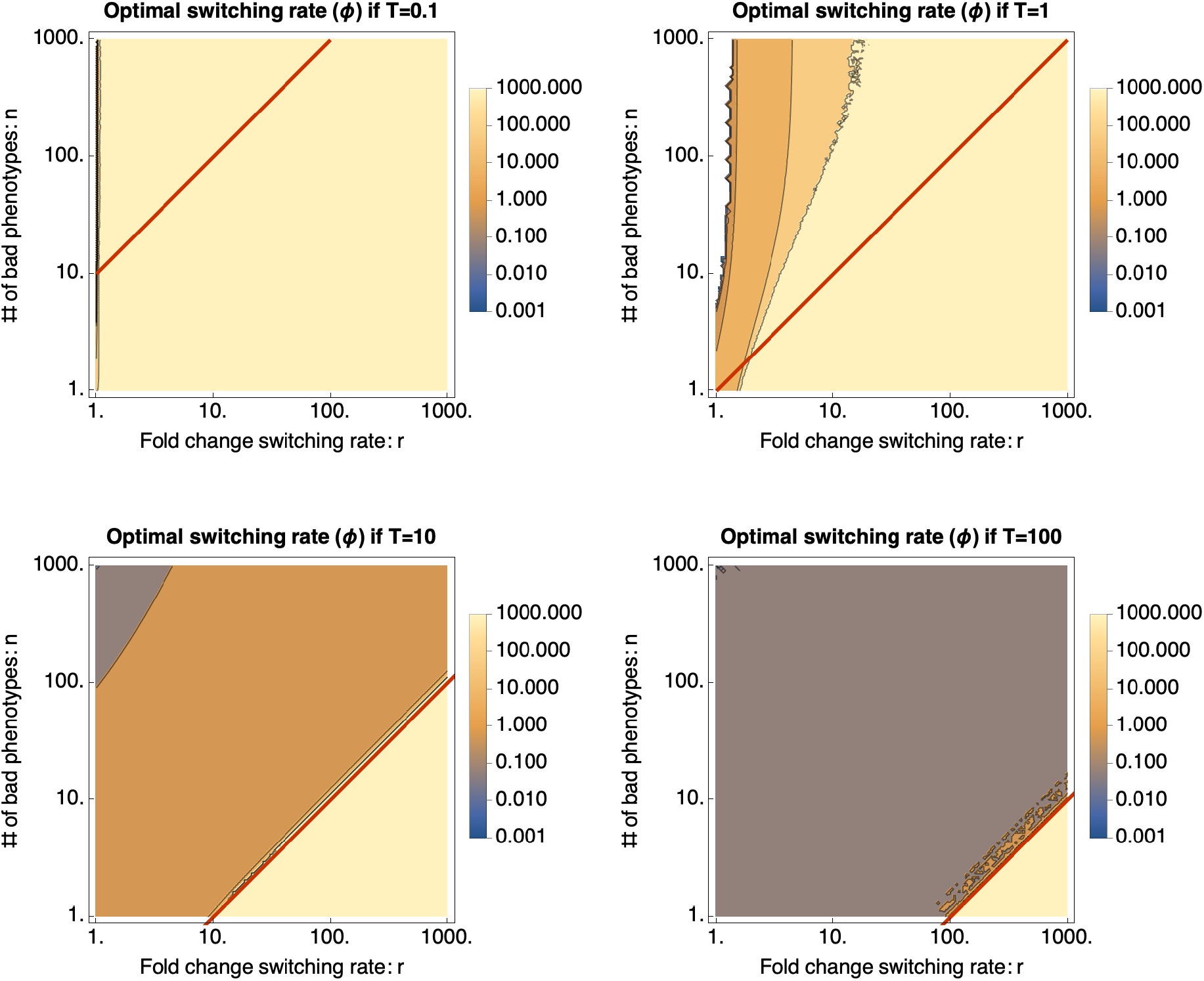
Optimal switching rate *ϕ* for a bet-hedging population that experiences GRDS. On the horizontal axes, we change the strength of GRDS, i.e. the ratio *r* of switching rates of slow and fast growing phenotypes. On the vertical axes, we change the number of bad phenotypes *n*. Each panel corresponds to a different environment duration T. The red line shows *T* = *r/n* which, for large enough *T*, divides the parameter regimes where *ϕ* ≈ 1/*T* (for *T* > *r/n*) and where *ϕ* → ∞ (for *T* < *r/n*). This figure was made with the Mathematica notebook `GRDS_minimal_model_analysis.nb`.

**Figure S7.**
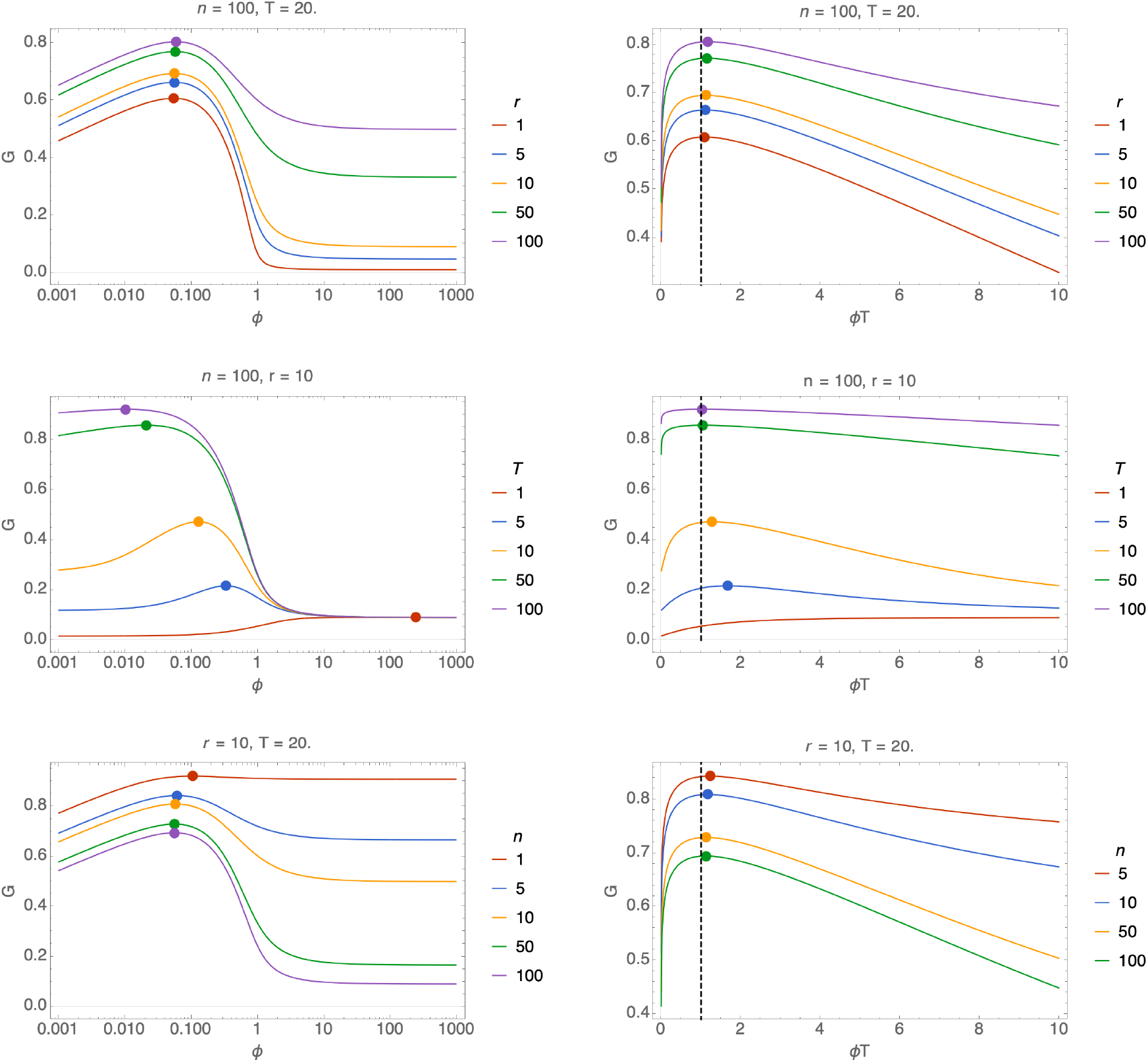
The optimal switching rate is approximately 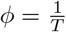. The left panels show the average population growth rate as a function of *ϕ* and the right panels as a function of *ϕT*. Other parameters are as indicated at the top of each panel and in the legends. The dots indicate the optimal switching rate and the dashed lines in the right panels show 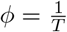. The optimal switching rates converge to 1/*T* when *T* becomes large (seen middle row of panels). This figure was made with the Mathematica notebook `GRDS_minimal_model_analysis.nb`.

**Figure S8.**
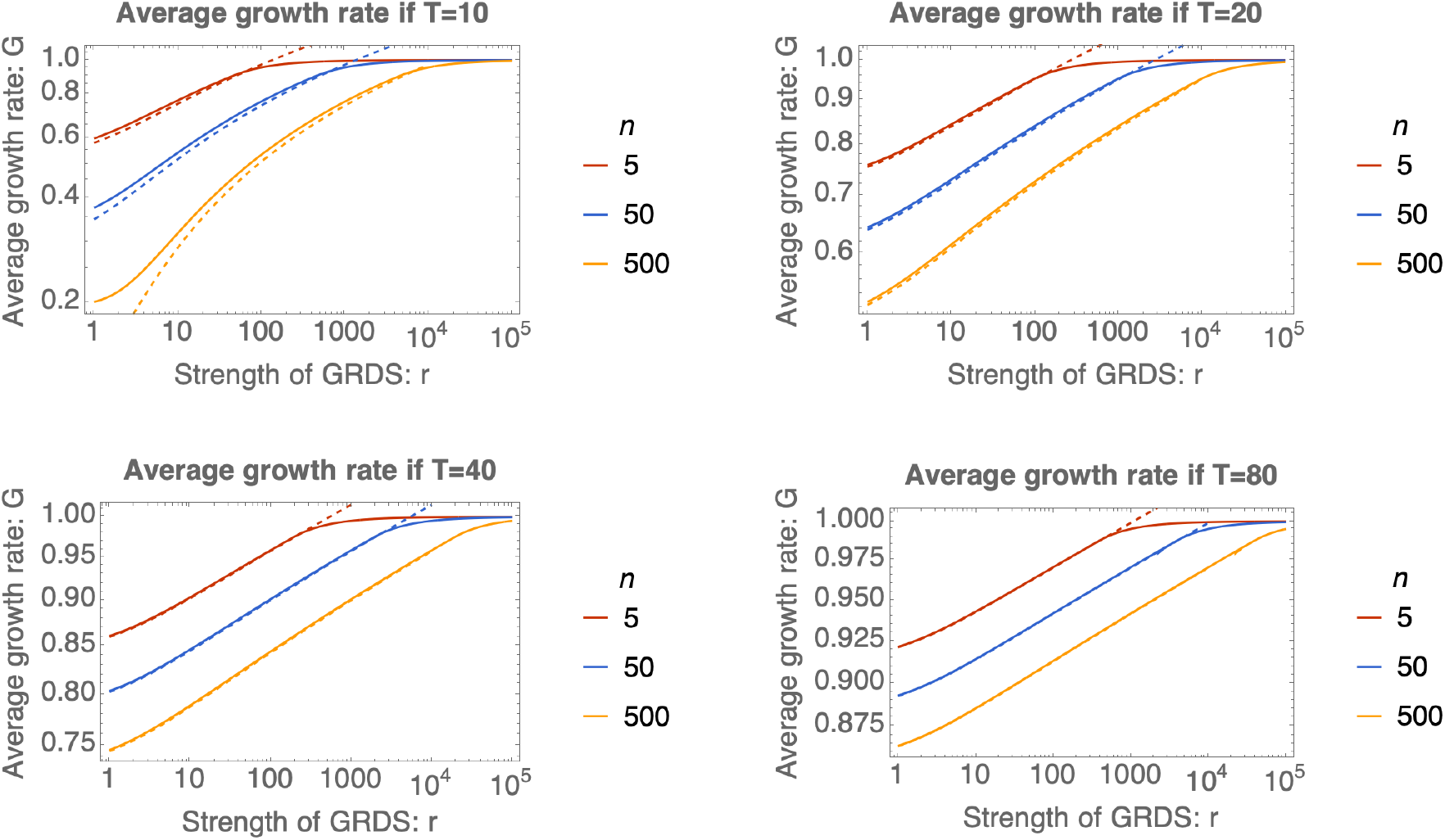
Analytical approximation of the average growth rate. Each panel shows *G* as a function of the strength of GRDS *r* (solid lines) and the analytical approximation 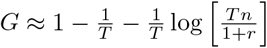, derived in Section 2 (dashed lines). Each panel corresponds to a different environment duration *T* (see titles), and each color corresponds to a different number of bad phenotypes *n* (see legends). This figure was made with the Mathematica notebook `GRDS_minimal_model_analysis.nb`.

### 2 Analytical derivations for the long-time limit of the general model

We will here first describe the original phenotype switching model of Kussell and Leibler [18]. We will describe the model formalism and their analytical approximation of the optimal solution. To keep this document self-contained, we will repeat the steps in the derivations by Kussell and Leibler and often add details where we think they are needed.

We will then extend this model with growth rate dependent switching rates. We will prove that growth rate-dependent switching increases a population’s average growth rate, in the situations where certain assumptions hold. These assumptions are clearly indicated in the text.

In this part of the SI dedicated to the general model, we will use *n* as the number of phenotypes instead of *n* +1, since in the general model there is not necessarily one phenotype that is adapted while the rest is not.

#### 2.A Prerequisite theory: Phenotype switching with constant switching rates

##### 2.A.1 The original phenotype switching model

The essential ingredients of the model are:

- *m* is the number of environments, *n* the number of phenotypes,
- 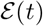 denotes a stochastic process determining which environment is active at time *t*
- the duration of an occurrence of environment *i* is drawn from an exponential distribution with mean *τ_I_*,
- the order in which the environments occur is determined by a Markov chain. The probability that environment *I* follows environment *J* is denoted by *b_IJ_*. Using these parameters, we can find the probability that environment *I* occurs as *p_I_*,
- a cell in phenotype i grows in environment *I* with rate 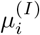,
- the switching rate from phenotype *j* to *i* in environment *I* is denoted by 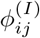.

Given these ingredients, the growth of the population can be modelled by the differential equation

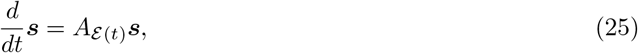

where 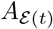 is the matrix belonging to the environment 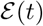. This matrix can be written as a sum of a diagonal matrix with the growth rates of all phenotypes in that environment, and a switching matrix Φ. The off-diagonal entries of the switching matrix are all positive, being filled with the abovementioned switching rates: 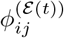. The diagonal elements are the negative sums of the corresponding columns, so that all cells that switch to a different phenotype are subtracted from the number of cells in their phenotype.

The total number of cells at time *t* is given by *N*(*t*) = ∑_*i*_ *s_i_*(*t*). We are interested in the average growth rate of the population over a long sequence of environments. We therefore define the long-term average growth rate as

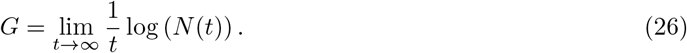

We use *G* instead of the Λ that was used in [18] to keep notation consistent across this work.

##### 2.A.2 Derivation of an analytic approximation for the average growth rate

In this subsection we will describe how the following approximation for the average growth rate can be derived

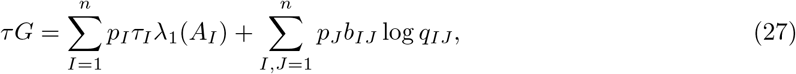

where *τ* is the average duration of an environment, and λ_1_(*A_I_*) is the dominant eigenvalue of the matrix *A_I_* corresponding to environment *I*. The terms *q_ij_* are obtained by projecting the dominant eigenvector of environment *J* onto the dominant eigenvector of environment *I*. The term log *q_IJ_* quantifies the loss of fitness after a transition in the environment due to the fraction of the population that is not in the optimal phenotype to grow in the new environment.

###### Slicing up time

The first step in the derivation is dividing the simulation time in intervals in which the environment does not change. This facilitates separating the different environments, which is practical since the model properties are markedly different in different environments. The length of these intervals are denoted *T_l_* and the cumulative time is denoted 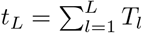. The state of the environment during the *l*-th interval is denoted *ϵ*(*l*).

###### Applying the theorem of Perron-Frobenius

The derivation depends on the assumption that the dominant eigenvalue of the matrices *A_I_* is real and that there is a gap between the dominant eigenvalue and the second eigenvalue. We can apply the Perron-Frobenius theorem to ensure these assumptions are met.

The Perron-Frobenius theorem can be applied only to nonnegative matrices, so that we will apply it to *A* + λ*I*, where *I* is the identity matrix. If λ is chosen large enough, this will give a matrix with only non-negative entries, because all off-diagonal elements of the switching matrix Φ are non-negative, and the growth rates only occur on the diagonal. When this matrix is in addition *primitive* we can apply the Perron-Frobenius theorem.

The matrix *A* + λ*I* is primitive when it is irreducible and has period 1. It is irreducible when any linear subspace spanned by standard basis vectors *e*_*i*1_,..., *e_ik_* with 0 < *i_k_* < *m* is not mapped into itself by *A* + λ*I*. This means that *A* + λ*I* is irreducible when no subset of phenotypes switches only to each other. This is generally the case for the system under consideration. *A* + λ*I* certainly has period 1 when the diagonal elements are all positive, which they are. Concluding, we generally have a nonnegative, primitive matrix *A* + λ*I*. We can thus apply the Perron-Frobenius theorem.

This implies that the dominant eigenvalue of *A* + λ*I* is a positive real number, *r*, which has a one-dimensional eigenspace. The corresponding right and left eigenvector have only positive entries. Note that *A* thus has dominant eigenvalue *r* – λ.

###### The eigenvectors in a single environment

For a given environment, we can choose the generalized eigenvectors 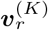 such that

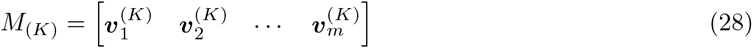

satisfies

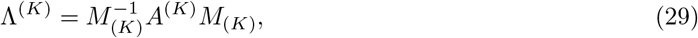

where Λ^(*K*)^ is in Jordan-block diagonal form with the eigenvalues, 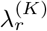, in decreasing order as coefficients.

###### Assuming distribution is stationary before environment switch

If the environment times, *T_l_*, are large enough, we can assume that the distribution of phenotypes at the switching time, *t*_*l*–1_, is equal to the distribution in the dominant eigenvector for this environment. So,

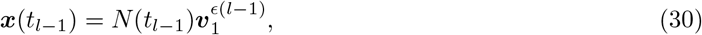

where *N*(*t*_*l*–1_) denotes the number of cells at time *t*_*l*–1_, and where we assume that the coefficients of the dominant eigenvectors, 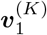, sum up to 1. In other words, we assume that the environments change slow enough to allow relaxation of the population to their stationary distribution.

###### Calculating time evolution in one environment

Starting from ***x***(*t*_*l*–1_), how will the vector evolve in environment *ϵ*(*l*)? We can find out by decomposing (projecting) ***x***(*t*_*l*–1_) in the generalized eigenvectors of *A*^(*ϵ*(*l*))^, which were grouped in *M*^(*ϵ*(*l*))^. The coordinates of ***x***(*t*_*l*–1_) in the new eigenbasis can be calculated with the inverse of the matrix of generalized eigenvectors:

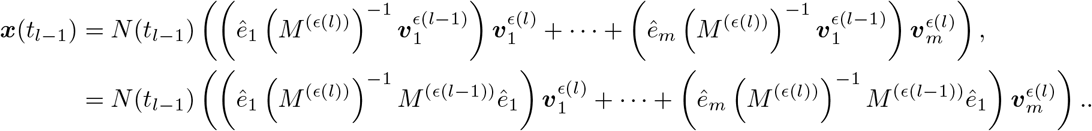

Note that this is just a sum of the generalized eigenvectors, weighted by some complicated coefficients. In case that we have a diagonalizable *A*, we can just multiply this by the time evolution matrix *A*^(*ϵ*(*l*))^. This will lead to each term being multiplied by the eigenvalue corresponding to the eigenvector in that term.

Although the initial weights for the other eigenvectors can be much larger, for large enough times the eigenvector 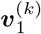 will again dominate the behaviour of ***x***(*t*), because it has the largest eigenvalue. Therefore, the growth of the population can be approximated by only considering this eigenvector. The coordinate of the new dominant eigenvector in the decomposition of the previous dominant eigenvectors in the new eigenvectors is denoted by

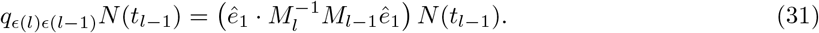

The time evolution of the number of cells in the *l*-th environment is thus given by

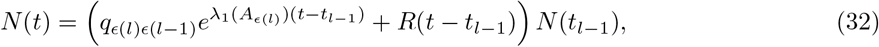

where *R*(*T*) is a function that grows slower than *e*^λ_1_ *A*_*ϵ*(*l*)_*T*^, and *R*(0) = 1 – *q*_*ϵ*(*l*)*ϵ*(*l*–1)_.

###### Composing the average growth rate expression

We can now put together the contributions of the separate environments to describe the growth of the population over the whole sequence of environments. From this, we can derive an expression for the average growth rate:

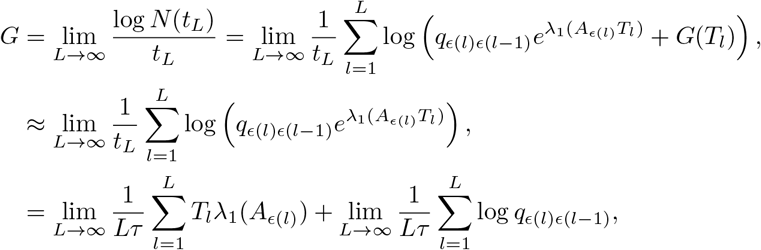

where 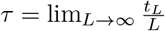 is the average duration of environments.

###### Validating the approximation

For the above derivation, we made the approximation that upon an environment switch, the system instantaneously projects to the dominant eigenvector of the new environment, and that all the other cells are lost. This might seem unreasonable but does not have large effects when the average durations of environments are relatively long. To be precise, we need

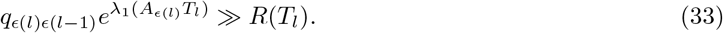

Now *R*(*T_l_*) can be overestimated when all eigenvalues are simple (i.e. each eigenvalue occurs only once). We get

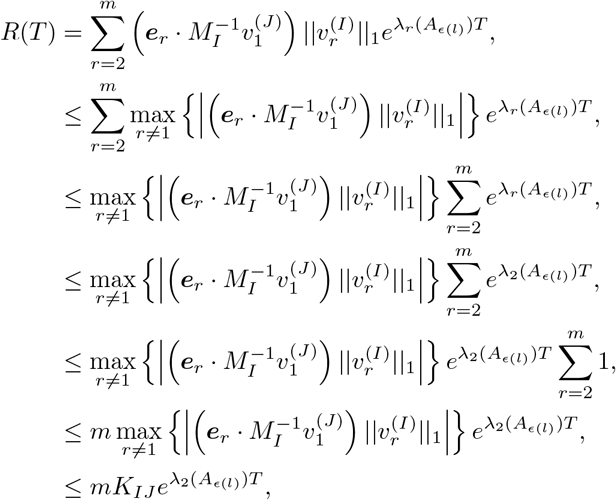

where *K_ij_* is defined appropriately. Combining this with the requirement (33) above, we get

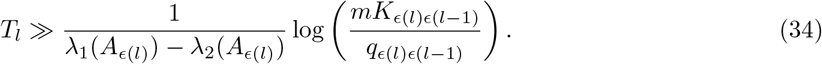

This approximation must hold for all switches between environments. Therefore, we take the maximum to get the requirement

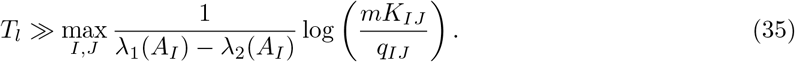

Importantly, note that the requirement is more easily met when the spectral gaps (λ_1_(*A_I_*) – λ_2_(*A_I_*)) are large.

###### Using the Markov chain to capture the order of environments

It was assumed that the environments change according to a Markov chain defined by switching rates *b_IJ_*. The probability of having environment *I* is denoted by *p_I_*. The duration of the *k*-th occurrence of environment *I* can be described by a random variable 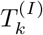, and it is assumed that the variables 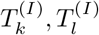 are independent, identically distributed variables with mean *τ_I_*.

For large *L*, the number of occurrences of environment *I* approaches *p_I_L*, and the number of subsequent occurrences of *J* and *I* approaches *p_J_b_IJ_L* Inserting this in the formula for the Lyapunov exponent gives

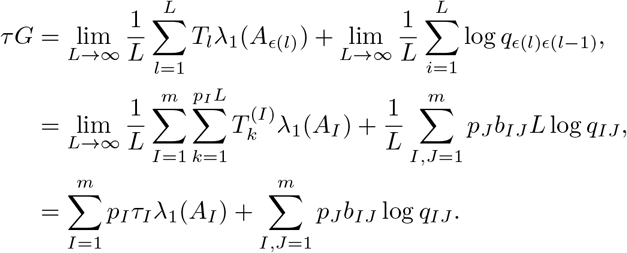

This completes the derivation of the expression in (27) of the average growth rate of the population.

#### 2.A.3 An approximation of the average growth rate in terms of growth and switching rates

To obtain analytical results about the effect of growth rate dependent switching rates on a population’s average growth rate, we need an expression of this average growth rate in terms of the switching rates. Starting with (27), we thus need to express λ_1_(*A_I_*) and log *q_IJ_* in terms of the model parameters. Both are determined by the eigenvalues and eigenvectors of the time evolution matrices *A_I_*. Following Kussell and Leibler [18], we find expressions for these eigenvalues and eigenvectors by using perturbation theory. This perturbation theory heavily relies on the assumption that the switching rates 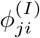 are small compared to the growth rates 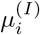, and we will see that we must even demand that the switching rates are small compared to the *growth rate differences.*

##### Using perturbation theory to obtain expressions for the eigenvalues and -vectors

We start by separating the matrix in two parts:

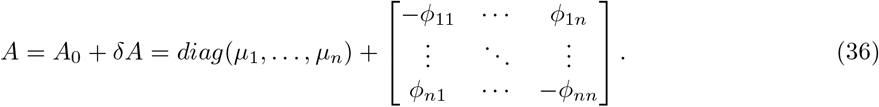

It is easy to find the eigenvalues and -vectors of *A*_0_, being:

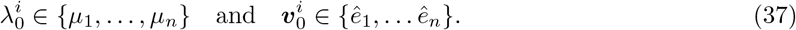

The question that we want to answer now is: Can we find *δ*λ^*i*^, *δ**v***, such that the set of eigenvalues 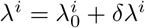, and eigenvectors 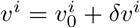 satisfies

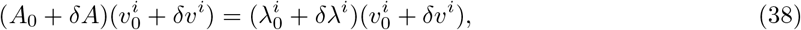

up to first order? We can expand this to get

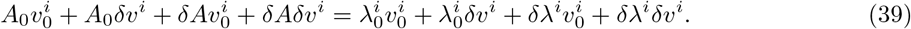

Then, we use that 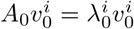, and we ignore all terms of order two and higher to get:

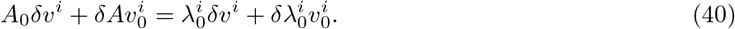

Since the unperturbed eigenvectors are just the elementary basis vectors, we can express the first order corrections to the eigenvectors in terms of the unperturbed ones:

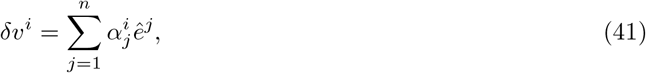

We insert the above decomposition into (40) and get

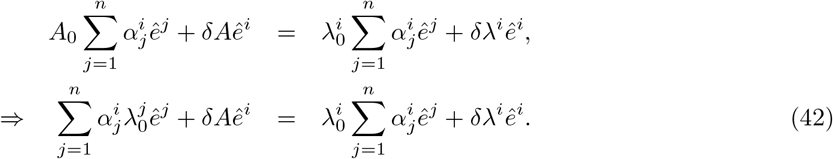

This is a set of *n* equalities, one for each choice of *i*. When we left-multiply this equation on both sides with the elementary unit vector *ê^k^*, we get

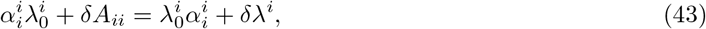

showing that we need *δ*λ^*i*^ = *δA_ii_* = ∑_*j*≠*i*_ *ϕ_ji_*. So, the first order perturbation decreases all eigenvalues by the sum of the ’away’-switching rates.

Then, we left-multiply (42) by *ê^k^* to get

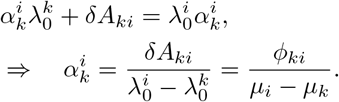

The only coefficients that are now undetermined are 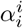. We can determine them by demanding that the eigenvectors are normalized. This yields:

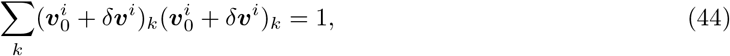

such that we get (to first order)

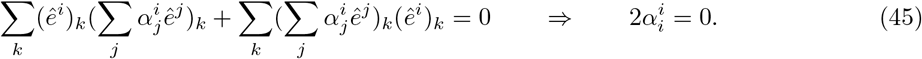

We thus get:

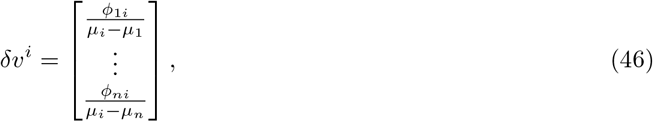

where the *i*-th coefficient is zero.

Concluding, the first order approximation of the eigenvalues are

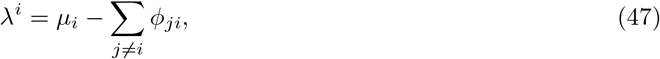

and of the eigenvectors:

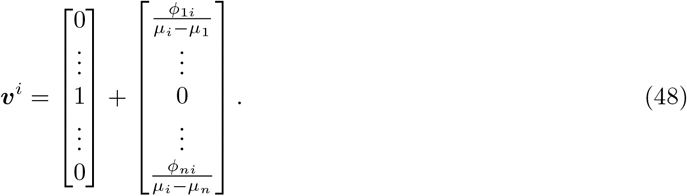

As described above, these eigenvectors will be collected in a square matrix

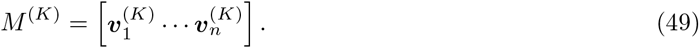

Note that the corrections to the eigenvectors only become small when the switching rates, *ϕ_ij_*, are small compared to the *differences* between growth rates, *μ_i_* – *μ_j_*.

##### Approximating the inverse of the eigenvector matrix

To obtain analytical approximations for the fractions *q_IJ_* in (27), we need to express the inverse of *M*^(*K*)^ in terms of the model parameters. Let the entries of this inverse matrix be given up to first order by 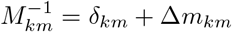. Now, we demand that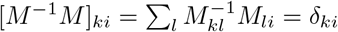. Expanding this expression gives

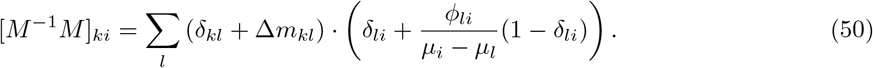

In case *k* = *i*, we get

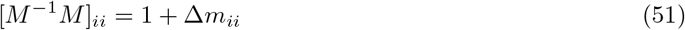

This implies that

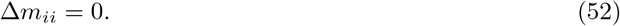

In the case that *k* ≠ *i* we get

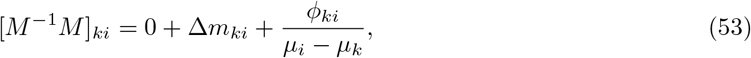

which implies

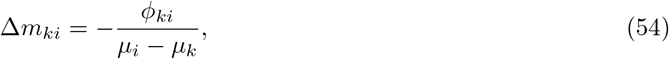

Concluding, the entries of (*M*^(*j*)^)^−1^ are

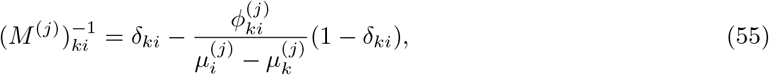

while

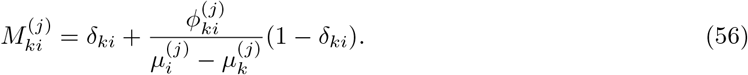

##### An expression for the projection of dominant eigenvectors

The fraction *q_IJ_* is obtained by projection of the dominant eigenvector in environment *J* on the dominant eigenvector of environment *I*. If we denote by *α_I_* the index of the dominant eigenvector in environment *I*, we get

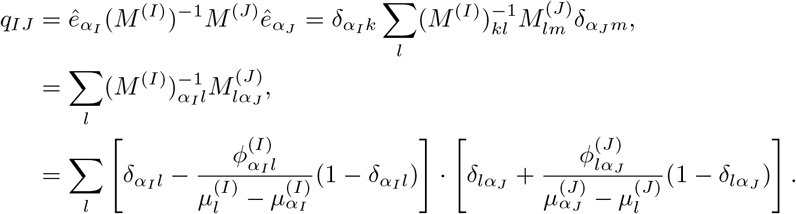

Expanding this multiplication gives if *α_I_* = *α_J_*:

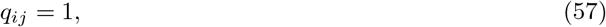

and if *α_I_* ≠ *α_J_*:

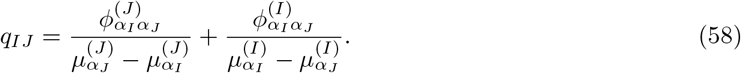

##### The average growth rate in terms of model parameters

Recall from Equation (27) that

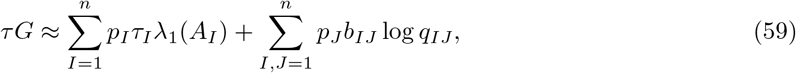

and note that we can insert the found expressions for λ_1_(*A_I_*) and *q_IJ_*. We get:

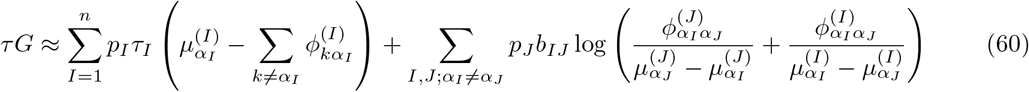

#### 2.A.4 The optimal constant switching rates

In [18], the switching rates were assumed independent of the current environment, just depending on the phenotypes the cell is switching between: 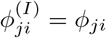. Also, to facilitate clearer notation they assume that the number of environments equals the number of phenotypes and that *α_i_* = *i*. We will repeat that last assumption here, although our results are true more generally. Using the analytical approximation for the average population growth rate in (60), we can find the optimal switching rates by setting the derivative equal to zero. This yields:

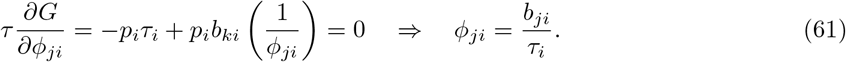

Inserting this in the formula for the average growth rate gives

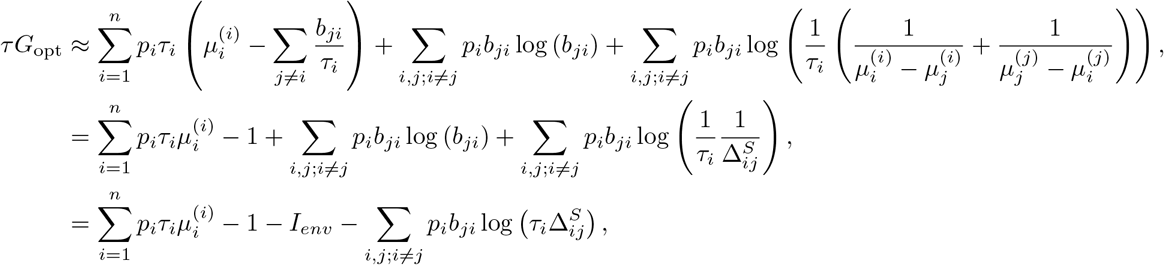

where 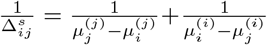, and 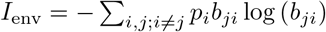 which is interpreted as the information entropy of the fluctuating environment.

#### 2.A.5 Interpretation of the constant switching results: explaining the trade-off between fast growth and fast adaptation

Let us consider the different terms in the expression for the average growth rate:

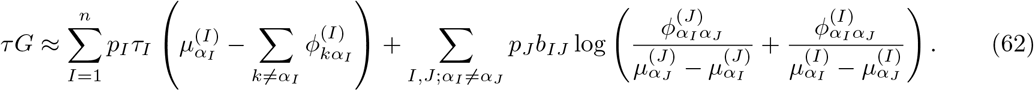

The first sum calculates an average, weighted by the occurrence of the environment, of the stationary growth rates in the different environments. In each environment the population growth rate is approximated by the growth rate of the fastest growth phenotype, minus the sum of the switching rates away from this phenotype. Through this term high switching rates away from fast growing phenotypes will thus negatively affect the average growth rate of the population.

The second sum of the term quantifies the loss in the average growth rate due to cells that are not instantaneously adapted to a new environment. Note that this term is negative because the switching rates in the numerators are generally much smaller than the growth rate differences in the denominators. The two terms in the logarithm show the two most important ways in which cells can adapt to a new environment by random switching. The first term shows that cells can have switched from the old to the new fastest growth phenotype before the environment switch. Through this term, high switching rates away from the fast growth phenotype will positively affect the average growth rate of the population. The second term in the logarithm shows that cells can switch from the old dominant phenotype to the new dominant phenotype after the switch. This shows that high switching rates from slow growth phenotypes to fast growth phenotypes can also benefit the average population growth rate.

Concluding, there is a clear trade-off between losing fitness due to a lower stationary growth rates at high switching rates, and losing fitness due to a longer adaptation time at low switching rates.

#### 2.A.6 List of assumptions

Our analytical investigation of random phenotype switching relies on several assumptions:

1. The duration of environments is long compared to division times and switching times
2. The spectral gaps (the difference between the first and second eigenvalue) of the time evolution matrices are large compared to the reciprocal of the average environment duration
3. The switching rates are small compared to the growth rates
4. The switching rates are small compared to the differences between the fastest growth rate and other growth rates in an environment

Note that we found that the optimal switching rates were found to be *ϕ_ji_* = *b_ji_/τ_i_*, so that assumptions 3 and 4 are expected to be satisfied automatically when the environment durations are long and the switching rates are optimised.

### 2.B Introducing growth rate dependent stability

In this section we will investigate if, and by how much, growth rate dependent phenotype switching increases the long-term growth rate of a population.

#### 2.B.1 Long-term population growth rates can be increased by growth rate dependency

First, we prove that we can always increase the long-term population growth rate when phenotype switching rates can be made dependent on the growth rates: 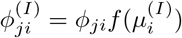.

##### Theorem 1.

*We consider population growth in a sequence of environments that satisfies the assumptions from Section 2.A.6. We start from a population with phenotype switching rates that are independent of the current environment. The long-term population growth rate can be increased by allowing for a dependency on the phenotype’s growth rate of the switching rate away from that phenotype.*

*Proof.* Let us take the expression for the average population growth rate as the benchmark:

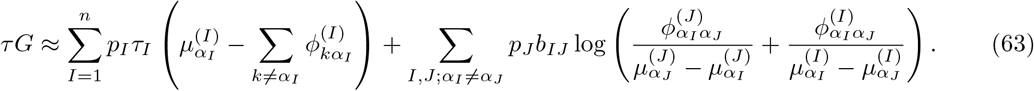

We introduce a growth rate dependence of the switching rates by assuming that the switching rates from non-dominant phenotypes increase 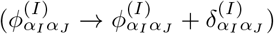, while switching rates from dominant phenotypes decrease 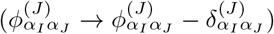. This gives

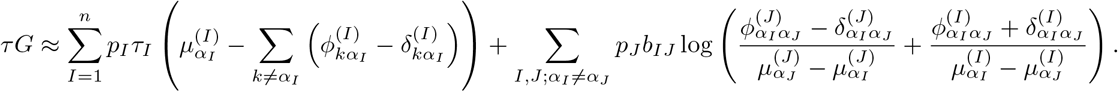

It is clear that the decrease of the switching rates away from optimal phenotypes provides a fitness benefit in the first term. This effect captures that fewer cells switch from the optimal phenotype in each environment, so that the stationary population growth rate is higher. Still, this beneficial effect might be undone by the changes in the terms of the second sum. We can show, however, that the overall change in the second sum is also positive, if the growth rate dependence is chosen well.

We define 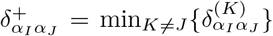, and 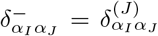, so that 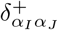 is the minimal increase in switching rate over all environments in which phenotype *α_J_* is not optimal. We then get

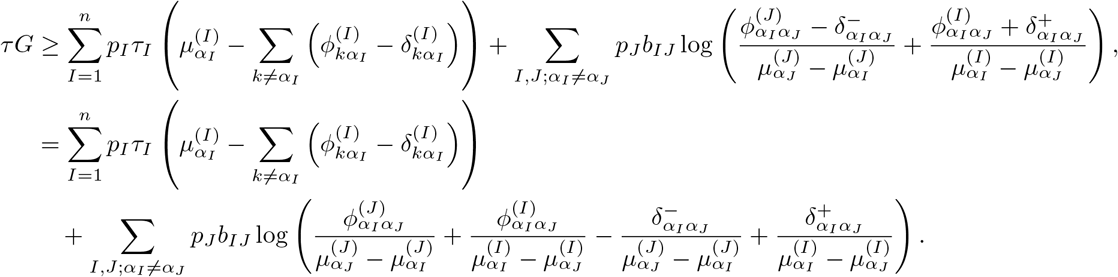

The growth rate dependence can now be chosen such that 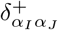 and 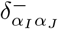 satisfy

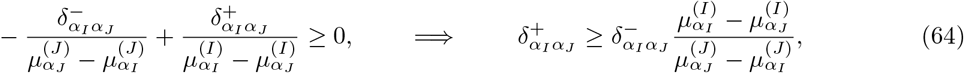

which implies that both terms in the expression for *τG* increase as a consequence of the growth rate dependence, which completes our proof.

We emphasise that the theorem was proven for any set of constant phenotype switching rates. This means that it does not matter if these phenotype switching rates were optimised or not, with growth rate dependent switching rates a population can always do better.

#### 2.B.2 Analytical approximation of the benefit of growth rate dependence

Consider a model of the type discussed in Section 2.A. We would like to get some estimate of how large the growth rate benefit due to growth rate dependent switching can become. To facilitate the analytical approximation, we will investigate the effects of this growth rate dependence in a simplified case. This will give us an expression that can be further simplified into the long-time limit of the minimal model described in Section 1. More general cases will be analysed numerically (Section 3). Here, we assume that:

1. the number of environments equals the number of phenotypes; each phenotype is optimal in one environment, and we will assume *α_i_* = *i* without loss of generality
2. the dominant phenotypes all have growth rate *μ*_max_,
3. the non-dominant phenotypes all have growth rate 0,
4. the factor difference between a switching rate from a slow-growth phenotype, and a switching rate from a fast-growth phenotype, is bounded by a parameter *r*.

Since we have only two possible growth rates, there are only two switching rates per pair of phenotypes: a switching rate away from fast-growth phenotypes 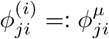, and a switching rate away from slow-growth phenotypes 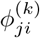, with the constraint that 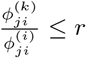.

We start by inserting these simplifying assumptions in the expression for the average population growth rate

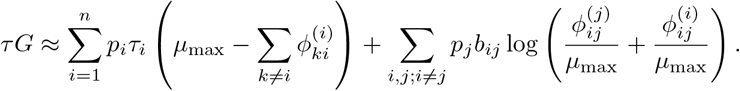

This expression shows that if the average growth rate is to be maximised, the switching rates away from slow-growth phenotypes 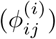 should be as high as possible. Therefore, we know that the constraint bounding the factor difference between high and low switching rates is always saturated in the optimum. As a consequence, we can write: 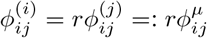, which gives

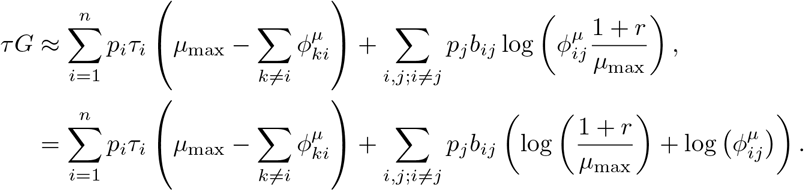

We can then find an analytical expression for the optimal switching rates by setting the derivative with respect to the switching rates to zero:

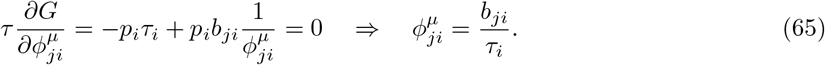

Note that these are the same optimal switching rates as in the non-growth rate dependent case. Inserting this in the formula for the average growth rate gives

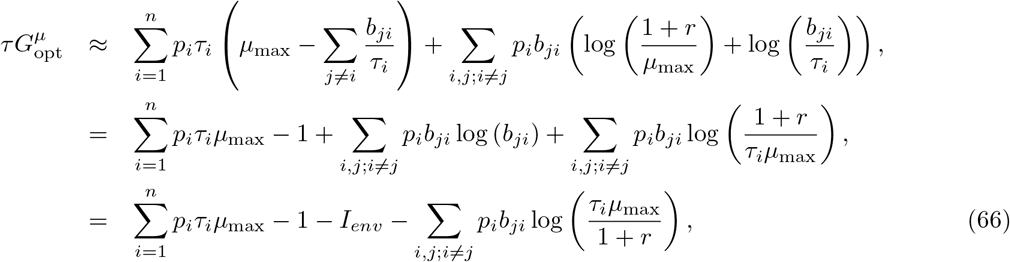

where 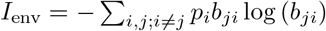 which is interpreted as the information entropy of the fluctuating environment.

We can compare this with the case where the switching rates were not growth rate dependent. In that case, and under the assumptions about the growth rates that we have used here, the expression for the long term growth rate is

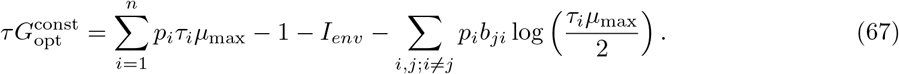

The growth rate benefit that growth rate dependent switching brings is thus

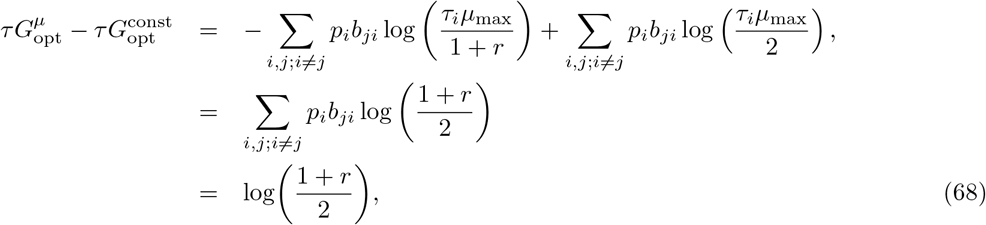

which implies that the average growth rate increases by

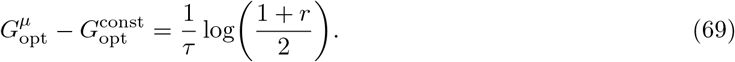

### 3 Numerics: Quantifying the advantage of growth rate dependent stability

We implemented the model described in Section 2.A in `Python`, which can be found at https://github.com/dhdegroot/GRDS-code-repository. In this section we will show the results of various numerical simulations to validate and elaborate on the intuition gained by the preceding analytical approach. We do this by performing many simulations for different choices of parameters to see if the fitness benefit of GRDS is present across all parameters, the results of which are shown in Figure 3 of the main text. We here explain which parameter sets were simulated.

For the first part of the parameters, we iterate their value over the elements from a list. All combinations of these parameter values were tested:

- the number of environments, *n*, was iterated over the set {4, 5, 8, 13, 19}
- the number of phenotypes, *m*, was either *n*/2, *n* or 2*n*; this number was rounded up when it was not an integer value.
- the average environment duration, *τ*, was picked from the set {1, 10.75, 20.5, 30.25, 40}. For each occurrence of an environment we sampled its duration from an exponential distribution with mean *τ*.
- for picking the growth rates of the various phenotypes in the different environments, we will need two parameters: the *spread* in growth rates of the fast phenotypes in different environments, *μ*_fast,spread_, and the minimal difference between a fast and a slow growth rate, *δ*. We iterated μfast,spread over {0.05, 0.2, 0.35, 0.5, 0.65}, and δ over {0.05, 0.1, 0.15, 0.2, 0.25}.

For the next subset of the parameters, each parameter value was drawn randomly for each of the combinations of the above parameters. We draw them in the following way:

- the Markov chain switching probabilities, *b_IJ_*, were drawn from a uniform distribution between 0 and 1, and then normalised such that ∑_*I*_ *b_IJ_* = 1.
- the maximal growth rates in the various environments, 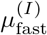, were drawn from a uniform distribution [0.9 − *μ*_fast,spread_, 0.9]
- the rest of the phenotypes in environment *I* have growth rates drawn from a uniform distribution on 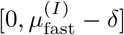.

Given these parameters, we now choose random switching rates:

- for the case of bet hedging, the constant phenotype switching rates *ϕ_ij_* were drawn uniformly in log-scale from the range: 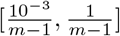. The dependence on the number of phenotypes *m* is introduced here to ensure that the maximal possible rate of switching out of a given phenotype is 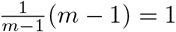.

Based on these switching rates, we then choose growth rate dependent switching rates, such that we can modify the strength of GRDS:

- for bet-hedging with GRDS of strength *r*, we take the range *R_ϕ_*(*r*) = [log(*ϕ_j_*) – 0.5 log(*r*), log(*ϕ_j_*) + 0.5 log(*r*)]. We subsequently determine the range of growth rates that cells in phenotype *j* can have in the different environments: 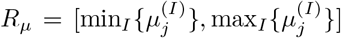, and we let *t^r^* be the linear map from *R_μ_* to *R_ϕ_*(*r*) that maps the minimal growth rate to the maximal switching rate and vice versa. The switching rate from phenotype *j* to *i* in environment *I* with GRDS of strength *r* is now determined by 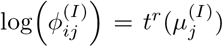. Based on the growth rate in the current environment, the switching rates between two phenotypes are thus interpolated in log-scale between an upper bound and a lower bound, where the factor difference between the upper and lower bound is *r*. To have a fair comparison between different strenghts of GRDS, we make sure that the average switching rate between two phenotypes 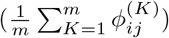 is the same for all strengths of GRDS. We do this by scaling all switching rates such that the average is equal to the originally drawn switching rate *ϕ_ij_*:

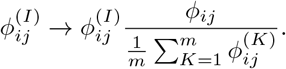

Given this complete set of parameters, we simulate population growth to compute the average population growth rate for various strengths of GRDS *r*. All simulations were done over a sequence of environments with a total duration that exceeded at least *n*^2^*τ*, so that we have had a reasonable sampling of the different environment switches that can occur.

## References

1. Opitz C, Ahrné E, Goldie KN, Schmidt A, Grzesiek S. Deuterium induces a distinctive *Escherichia coli* proteome that correlates with the reduction in growth rate. The Journal of biological chemistry. 2019 feb;294(7):2279–2292.

2. Donaldson-Matasci MC, Bergstrom CT, Lachmann M. The fitness value of information. Oikos. 2010 feb;119(2):219–230. Available from: https://doi.org/10.1111/j.1600-0706.2009.17781.x.

3. Rivoire O, Leibler S. The value of information for populations in varying environments. Journal of Statistical Physics. 2011;142(6):1124–1166. Available from: https://doi.org/10.1007/s10955-011-0166-2.

4. Berg HC, Purcell EM. Physics of chemoreception. Biophysical journal. 1977 nov;20(2):193–219.

5. Bialek W, Setayeshgar S. Physical limits to biochemical signaling. Proceedings of the National Academy of Sciences. 2005;102(29):10040–10045. Available from: https://www.pnas.org/doi/abs/10.1073/pnas.0504321102.

6. Dong H, Nilsson L, Kurland CG. Gratuitous overexpression of genes in *Escherichia coli* leads to growth inhibition and ribosome destruction. Journal of Bacteriology. 1995 mar;177(6):1497–1504. Available from: https://pubmed.ncbi.nlm.nih.gov/7883706/ http://www.ncbi.nlm.nih.gov/pubmed/7883706 http://www.pubmedcentral.nih.gov/articlerender.fcgi?artid=PMC176765.

7. Scott M, Gunderson CW, Mateescu EM, Zhang Z, Hwa T. Interdependence of cell growth and gene expression: origins and consequences. Science. 2010;330(6007):1099–1102. Available from: http://www.ncbi.nlm.nih.gov/pubmed/21097934.

8. Govern CC, ten Wolde PR. Optimal resource allocation in cellular sensing systems. Proceedings of the National Academy of Sciences. 2014 dec;111(49):17486LP–17491. Available from: http://www.pnas.org/content/111/49/17486.abstract.

9. Smits WK, Kuipers OP, Veening JW. Phenotypic variation in bacteria: The role of feedback regulation. Nature Reviews Microbiology. 2006 apr;4(4):259–271. Available from: http://www.nature.com/articles/nrmicro1381.

10. Thattai M, Van Oudenaarden A. Stochastic gene expression in fluctuating environments. Genetics. 2004 may;167(1):523–530. Available from: http://www.ncbi.nlm.nih.gov/pubmed/15166174 http://www.pubmedcentral.nih.gov/articlerender.fcgi?artid=PMC1470854.

11. Balaban NQ, Merrin J, Chait R, Kowalik L, Leibler S. Bacterial persistence as a phenotypic switch. Science. 2004 sep;305(5690):1622LP–1625. Available from: http://science.sciencemag.org/content/305/5690/1622.abstract.

12. Kærn M, Elston TC, Blake WJ, Collins JJ. Stochasticity in gene expression: From theories to phenotypes. Nature Reviews Genetics. 2005;6(6):451–464. Available from: https://pubmed.ncbi.nlm.nih.gov/15883588/.

13. Beaumont HJE, Gallie J, Kost C, Ferguson GC, Rainey PB. Experimental evolution of bet hedging. Nature. 2009;462(7269):90–93. Available from: http://dx.doi.org/10.1038/nature08504.

14. Simons AM. Modes of response to environmental change and the elusive empirical evidence for bet hedging. Proceedings of the Royal Society B: Biological Sciences. 2011;278(1712):1601–1609. Available from: https://pubmed.ncbi.nlm.nih.gov/21411456/.

15. Levy SF, Ziv N, Siegal ML. Bet hedging in yeast by heterogeneous, age-correlated expression of a stress protectant. PLoS Biology. 2012 may;10(5):1001325. Available from: www.plosbiology.org.

16. Lewontin CR, Cohen D. On population growth rate in a randomly varying environment. Proceedings of the National Academy of Sciences. 1969 apr;62(4):1056–1060. Available from: https://doi.org/10.1073/pnas.62.4.1056.

17. Karlin S, Lieberman U. Random temporal variation in selection intensities: case of large population size. Theoretical population biology. 1974 dec;6(3):355–382.

18. Kussell E, Leibler S. Phenotypic diversity, population growth, and information in fluctuating environments. Science. 2005;309(5743):2075–2078.

19. Gaál B, Pitchford JW, Wood AJ. Exact results for the evolution of stochastic switching in variable asymmetric environments. Genetics. 2010;184(4):1113–1119.

20. Acar M, Mettetal JT, Van Oudenaarden A. Stochastic switching as a survival strategy in fluctuating environments. Nature Genetics. 2008;40(4):471–475.

21. New AM, Cerulus B, Govers SK, Perez-samper G, Zhu B, Boogmans S, et al. Different levels of catabolite repression optimize growth in stable and variable environments. PLoS Biology. 2014;12(1):17–20.

22. Keren L, Van Dijk D, Weingarten-Gabbay S, Davidi D, Jona G, Weinberger A, et al. Noise in gene expression is coupled to growth rate. Genome Research. 2015;25(12):1893–1902. Available from: https://pubmed.ncbi.nlm.nih.gov/26355006/.

23. Kiviet DJ, Nghe P, Walker N, Boulineau S, Sunderlikova V, Tans SJ. Stochasticity of metabolism and growth at the single-cell level. Nature. 2014 oct;514(7522):376–379.

24. Urchueguía A, Galbusera L, Chauvin D, Bellement G, Julou T, van Nimwegen E. Genome-wide gene expression noise in *Escherichia coli* is condition-dependent and determined by propagation of noise through the regulatory network. PLoS Biology. 2021 dec;19(12):e3001491. Available from: https://journals.plos.org/plosbiology/article?id=10.1371/journal.pbio.3001491.

25. Biggar SR, Crabtree GR. Cell signaling can direct either binary or graded transcriptional responses. EMBO Journal. 2001;20(12):3167–3176. Available from: https://pubmed.ncbi.nlm.nih.gov/11406593/.

26. Thattai M, Shraiman BI. Metabolic switching in the sugar phosphotransferase system of *Escherichia coli*. Biophysical Journal. 2003;85(2):744–754. Available from: https://pubmed.ncbi.nlm.nih.gov/12885625/.

27. Norman TM, Lord ND, Paulsson J, Losick R. Stochastic switching of cell fate in microbes. Annual Review of Microbiology. 2015;69(1):381–403. Available from: http://www.ncbi.nlm.nih.gov/pubmed/26332088.

28. Raj A, van Oudenaarden A. Nature, nurture, or chance: stochastic gene expression and its consequences. Cell; 2008. Available from: http://www.ncbi.nlm.nih.gov/pubmed/18957198 http://www.pubmedcentral.nih.gov/articlerender.fcgi?artid=PMC3118044.

29. Eldar A, Elowitz MB. Functional roles for noise in genetic circuits. Nature. 2010 sep;467(7312):167–173. Available from: http://www.ncbi.nlm.nih.gov/pubmed/20829787 http://www.pubmedcentral.nih.gov/articlerender.fcgi?artid=PMC4100692.

30. Kashiwagi A, Urabe I, Kaneko K, Yomo T. Adaptive response of a gene network to environmental changes by fitness-induced attractor selection. PLoS ONE. 2006;1(1).

31. Furusawa C, Kaneko K. A generic mechanism for adaptive growth rate regulation. PLoS Computational Biology. 2008;4(1):0035–0042.

32. King OD, Masel J. The evolution of bet-hedging adaptations to rare scenarios. Theoretical Population Biology. 2007 dec;72(4):560–575. Available from: https://pubmed.ncbi.nlm.nih.gov/17915273/.

33. Kelly JL. A New Interpretation of Information Rate. Bell System Technical Journal. 1956 jul;35(4):917–926. Available from: https://ieeexplore.ieee.org/document/6771227.

34. Slatkin M. Hedging one’s evolutionary bets. Nature. 1974;250(5469):704–705. Available from: https://doi.org/10.1038/250704b0.

35. Ozbudak EM, Thattal M, Lim HH, Shraiman BI, Van Oudenaarden A. Multistability in the lactose utilization network of *Escherichia coli*. Nature. 2004 feb;427(6976):737–740. Available from: https://www.nature.com/articles/nature02298.

36. Maamar H, Dubnau D. Bistability in the *Bacillus subtilis* K-state (competence) system requires a positive feedback loop. Molecular microbiology. 2005 may;56(3):615–624. Available from: https://pubmed.ncbi.nlm.nih.gov/15819619/.

37. Kearns DB, Losick R. Cell population heterogeneity during growth of *Bacillus subtilis*. Genes & development. 2005 dec;19(24):3083–3094. Available from: https://pubmed.ncbi.nlm.nih.gov/16357223/.

38. Rainey PB, Beaumont HJE, Ferguson GC, Gallie J, Kost C, Libby E, et al. The evolutionary emergence of stochastic phenotype switching in bacteria. Microbial Cell Factories. 2011 aug;10(SUPPL. 1):1–7. Available from: https://microbialcellfactories.biomedcentral.com/articles/10.1186/1475-2859-10-S1-S14.

39. Bull JJ. Evolution of phenotypic variance. Evolution. 1987;41(2):303–315. Available from: https://pubmed.ncbi.nlm.nih.gov/28568756/.

40. Haccou P, Iwasa Y. Optimal mixed strategies in stochastic environments. Theoretical Population Biology. 1995 apr;47(2):212–243.

41. Kopf SH, McGlynn SE, Green-Saxena A, Guan Y, Newman DK, Orphan VJ. Heavy water and 15N labelling with NanoSIMS analysis reveals growth rate-dependent metabolic heterogeneity in chemostats. Environmental Microbiology. 2015 jul;17(7):2542–2556. Available from: http://doi.wiley.com/10.1111/1462-2920.12752.

42. Schreiber F, Littmann S, Lavik G, Escrig S, Meibom A, Kuypers MMM, et al. Phenotypic heterogeneity driven by nutrient limitation promotes growth in fluctuating environments. Nature Microbiology. 2016 may;1(6):1–7. Available from: https://www.nature.com/articles/nmicrobiol201655.

43. Nikolic N, Schreiber F, Dal Co A, Kiviet DJ, Bergmiller T, Littmann S, et al. Cell-to-cell variation and specialization in sugar metabolism in clonal bacterial populations. PLoS Genetics. 2017;13(12):1–24.

44. Zimmermann M, Escrig S, Lavik G, Kuypers MMM, Meibom A, Ackermann M, et al. Substrate and electron donor limitation induce phenotypic heterogeneity in different metabolic activities in a green sulphur bacterium. Environmental Microbiology Reports. 2018 apr;10(2):179–183.

45. Schreiber F, Ackermann M. Environmental drivers of metabolic heterogeneity in clonal microbial populations. Elsevier Ltd; 2020.

46. Gasperotti A, Brameyer S, Fabiani F, Jung K. Phenotypic heterogeneity of microbial populations under nutrient limitation. Elsevier Ltd; 2020.

47. Julou T, Zweifel L, Blank D, Fiori A, van Nimwegen E. Subpopulations of sensorless bacteria drive fitness in fluctuating environments. PLoS Biology. 2020 dec;18(12):e3000952. Available from: https://doi.org/10.1371/journal.pbio.3000952.

48. Kleijn IT, Krah LHJ, Hermsen R. Noise propagation in an integrated model of bacterial gene expression and growth. PLoS Computational Biology. 2018;14(10):1–18.

49. Bertaux F, Marguerat S, Shahrezaei V. Division rate, cell size and proteome allocation: Impact on gene expression noise and implications for the dynamics of genetic circuits. Royal Society Open Science. 2018 mar;5(3):172234. Available from: https://royalsocietypublishing.org/doi/10.1098/rsos.172234.

50. Julou T, Gervais T, Blank D, van Nimwegen E. Growth rate controls the sensitivity of gene regulatory circuits. bioRxiv. 2022 jan;p. 2022.04.03.486858. Available from: http://biorxiv.org/content/early/2022/04/04/2022.04.03.486858.abstract.

51. Wolf L, Silander OK, van Nimwegen E. Expression noise facilitates the evolution of gene regulation. eLife. 2015;4(JUNE):1–48.

